# DNA damage-induced PARP/ALC1 activation leads to Epithelial-to-Mesenchymal transition stimulating homologous recombination

**DOI:** 10.1101/2024.01.16.575847

**Authors:** Fatemeh Rajabi, Rebecca Smith, Win-Yan Liu-Bordes, Michael Schertzer, Sebastien Huet, Arturo Londoño-Vallejo

**Affiliations:** Institut Curie, CNRS-UMR3244, Sorbonne University, 75005 Paris, France; Cancer Genomics lab, Inserm-U981, Gustave Roussy Cancer Center Grand Paris, Villejuif, 94805, France; Univ Rennes, CNRS, IGDR (Institut de génétique et développement de Rennes) - UMR 6290, BIOSIT – UMS3480, F- 35000 Rennes, France; Sir William Dunn School of Pathology, University of Oxford, South Parks Road, Oxford, OX1 3RE, UK; Institut Curie, Inserm U1021-CNRS UMR 3347, Paris Saclay University, Centre Universitaire, 91405 Orsay Cedex, France

**Keywords:** DNA induced-, PARP/ALC1 dependent-, EMT stimulates DNA repair through homologous recombination

## Abstract

Epithelial-to-mesenchymal transition (EMT) allows cancer cells to metastasize while acquiring resistance to apoptosis and to chemotherapeutic agents with significant implications in patients’ prognosis and survival. Despite its clinical relevance, the mechanisms initiating EMT during cancer progression remain poorly understood. We demonstrate that DNA damage triggers EMT by activating PARP and the PARP-dependent chromatin remodeler ALC1 (CHD1L). We show that this activation directly facilitates the access to chromatin of EMT transcriptional factors (TFs) which then initiate cell reprogramming. We also show that EMT-TFs bind to the RAD51 promoter to stimulate its expression and to promote DNA repair by recombination. Importantly, a clinically relevant PARP inhibitor totally reversed or prevented EMT in response to DNA damage while resensitizing tumor cells to other genotoxic agents. Overall, our observations shed light on the intricate relationship between EMT, DNA damage response and PARP inhibitors, providing valuable insights for future therapeutic strategies in cancer treatment.

## Introduction

The epithelial-to-mesenchymal transition (EMT) is a developmental process that sustains organogenesis. It enables embryonic progenitor epithelial cells to migrate and colonize new niches, followed by mesenchymal-to-epithelial transition (MET) and subsequent differentiation (reviewed in (*1*)). This phenomenon is governed by six transcriptional factors that trigger global changes in gene expression, thus driving major cytoskeleton reorganization and conferring cell mobility properties (*2*). In adult organisms, a similar program occurs during physiological healing processes and pathologically during tumor progression. EMT has been linked to the metastatic process, as well as with the acquisition of stem-like properties and resistance to chemotherapeutic agents by tumor cells (*3, 4*). Multiple pathways have been shown to modulate the EMT program in cancer cells but little is known about the mechanisms responsible to initiate EMT in the primary tumor. Previous work in our laboratory (*5, 6*) has shown that persistent endogenous DNA damage resulting from repeated telomere-driven breakage-fusion-bridge cycles promotes EMT in transformed human epithelial kidney (HEK) cells. This observation suggests that EMT may occur in early carcinoma cells exhibiting chromosome instability (CIN) (*7-9*). Notably, an EMT-compatible signature has been detected in experimental models of metastatic cancer in association with CIN (*10*).

CIN stands out as one of the most pervasive hallmarks of cancer cells (*11*). Direct evidence supporting its contribution to tumorigenesis has come from clinical observations associating inherited mutations in genes involved in genome maintenance and DNA repair with an elevated risk for carriers to develop carcinomas (*12-14*). A paradigmatic example of such an association is found in BRCA1/2 constitutive mutations, correlating with an increased susceptibility to breast and ovarian cancers (*15*). Furthermore, in diverse cancer types, including breast and ovarian cancers, there is a direct correlation between the degree of CIN in the primary tumor and the increased risk of death attributed to metastatic disease (*16, 17*). Despite the clinical significance of CIN in cancer progression, the molecular mechanisms linking CIN to the ability of tumors to metastasize in vivo are not fully understood. These connections likely encompass both cell-autonomous mechanisms, such as the inflammatory response triggered by the detection of double-stranded DNA in the cytoplasm (*10*), as well as cell non-autonomous ones, such as a local senescent or pro-inflammatory microenvironment (*6, 18, 19*).

An initial molecular link between the DNA damage response (DDR) and EMT was established by a prior study showing that loss of H2AX, a primary sensor of DNA damage, leads to EMT (*20*). While the study concluded that EMT occurred likely due to chromatin-mediated enhanced activity of EMT-related transcription factors (EMT-TFs), it did not elucidate the mechanistic basis of the DDR-EMT relationship. To address this gap, we directly explored this connection by examining the impact on EMT upon inactivation of other non-histone DDR components. Our findings reveal that disabling 53BP1 leads to spontaneous EMT in a variety of tumor cells and that DDR inactivation does not impede further EMT reinforcement in response to exogenous damage. Moreover, 53BP1-abrogated cells showed increased expression of RAD51 in an EMT-TF-dependent manner and increased resistance to drugs. Mechanistically, we show that inhibition of PARP or the abolition of ALC1 (CHD1L), a chromatin remodeler dependent on PARP activation, leads to total reversal of EMT-associated phenotypes in DDR-deficient cells and sensitizes them to other drugs. Intriguingly, Olaparib, a PARP1/2 inhibitor, completely reversed the DDR-related EMT and restored drug sensitivity of EMT tumor cells. This study advocates for the combination of PARP inhibitors with other classic genotoxic agents to mitigate the induction of EMT by the latter.

## Results

### Hallmarks of EMT are rapidly induced in response to DNA damage

We have previously shown that spontaneous CIN in mortal Human Epithelial Kidney cells as well as long-term exposure of immortal HEK cells to DNA damage agents like etoposide lead to EMT (*5*). To determine how soon after initial DNA damage EMT-related changes can be detected, we used the latter experimental system to precisely monitor in time responses such as cell morphology changes and gene expression of EMT-associated transcriptional factors (TFs), micro-RNAs (miRs) and other EMT-related markers. As early as 7 days after treatment initiation with low doses of etoposide (0.1 µM), epithelial cells displayed morphological changes, from cobble-like, tightly packed cells, to elongated, loosely packed fibroblastoid-like cells (Supplemental figure S1A). These changes were associated with a strong increase in the expression of all six EMT-TFs, as well as with an upregulation of mesenchymal markers including Vimentin, THBS1, MMP3, Seerpine1 and FN1, along with a significant downregulation of epithelial markers such as E-cadherin (Supplemental Figure S1B). Most of these changes could be also detected at the protein level (Supplemental figure S1C) including some EMT-TFs. In addition, etoposide-treated cells displayed a significant down-regulation in the EMT-related miRs miR-200a, miR-200b, miR-200c (Supplemental Figure S1D). To ensure that this phenomenon is not exclusive of kidney-derived cell lines, we also treated with low doses of etoposide MCF7 and HCT116 cancer cell lines, from breast and colon origin respectively, as well as the transformed prostate epithelial cell line PNT1A. As shown in Supplemental Figure S1E, all cell lines displayed a similar, albeit not identical, trend in expression changes affecting EMT-associated genes with downregulation of E-Cadherin in the case of MCF7 and HCT116, upregulation of some mesenchymal markers and of at least one EMT-TF in all cases. Altogether, these results demonstrate that DNA damage inflicted to epithelial cells rapidly induces an EMT response that can robustly be followed by cell morphology changes and changes in the expression of EMT-TFs and other EMT-associated markers.

### Blunting the DNA Damage Response propels transformed epithelial cells into EMT

To unravel the role of DDR in triggering EMT, we searched to curtail the pathway by inactivating key DNA damage mediators such as 53BP1. 53BP1 is also a major contributor to repair pathway choice in response to double strand breaks (DSB) by protecting these breaks from resection thus favoring non-homologous end joining (NHEJ) over homologous recombination (HR) (*21*). We used the Cas9 (D10A)-based approach described by Chiang et al (*22*) to transform, sort and clone HEK-Early cells. Two 53BP1-KO clones were isolated with a complete inactivation of the locus (Figure 1A). Unexpectedly, 53BP1-KO cell clones spontaneously displayed a mesenchymal-like morphology (Figure 1B) and a higher expression of EMT-TFs along with the mesenchymal markers MMP3, FN1, Vimentin, THBS1, and Serpine1 (Figure 1C-D). Epithelial markers, on the other hand, showed diminished mRNA expression levels in 53BP1-KO cells (Figure 1C). Finally, miRNA(miR)-200 family members, which are key regulatory factors to maintain epithelial identity, were significantly downregulated in KO cells with respect to parental cells (Figure 1E), in agreement with the observed loss of epithelial characteristics. These results are highly reminiscent of those made by Weyemi et al, after elimination of H2AX, also a DDR sensor. To ascertain that the EMT-related phenotypes detected in 53BP1-KO clones were not due to off-target effects, we conducted rescue experiments by exogenously expressing 53BP1. 53BP1-KO cells were transfected with pcDNA3.1-53BP1 and kept in batch under antibiotic selection. Upon expression of exogenous 53BP1, the morphological features of 53BP1-KO cells displayed a remarkable shift from spindle-like, isolated mesenchymal cells towards packed, cobblestone, epithelial cells (Figure 1F). Further analyses confirmed this reversal as they revealed a significant downregulation of EMT-TFs and other mesenchymal markers such as MMP3, FN1, along with an increase in the expression of the epithelial markers E-cadherin, Lumican, and CDH3 (Figure 1G-H). The fact that 53BP1-KO cells were able to undergo MET when 53BP1 expression was restored underscores the fundamental plasticity that characterizes all EMT.

**Figure 1.**
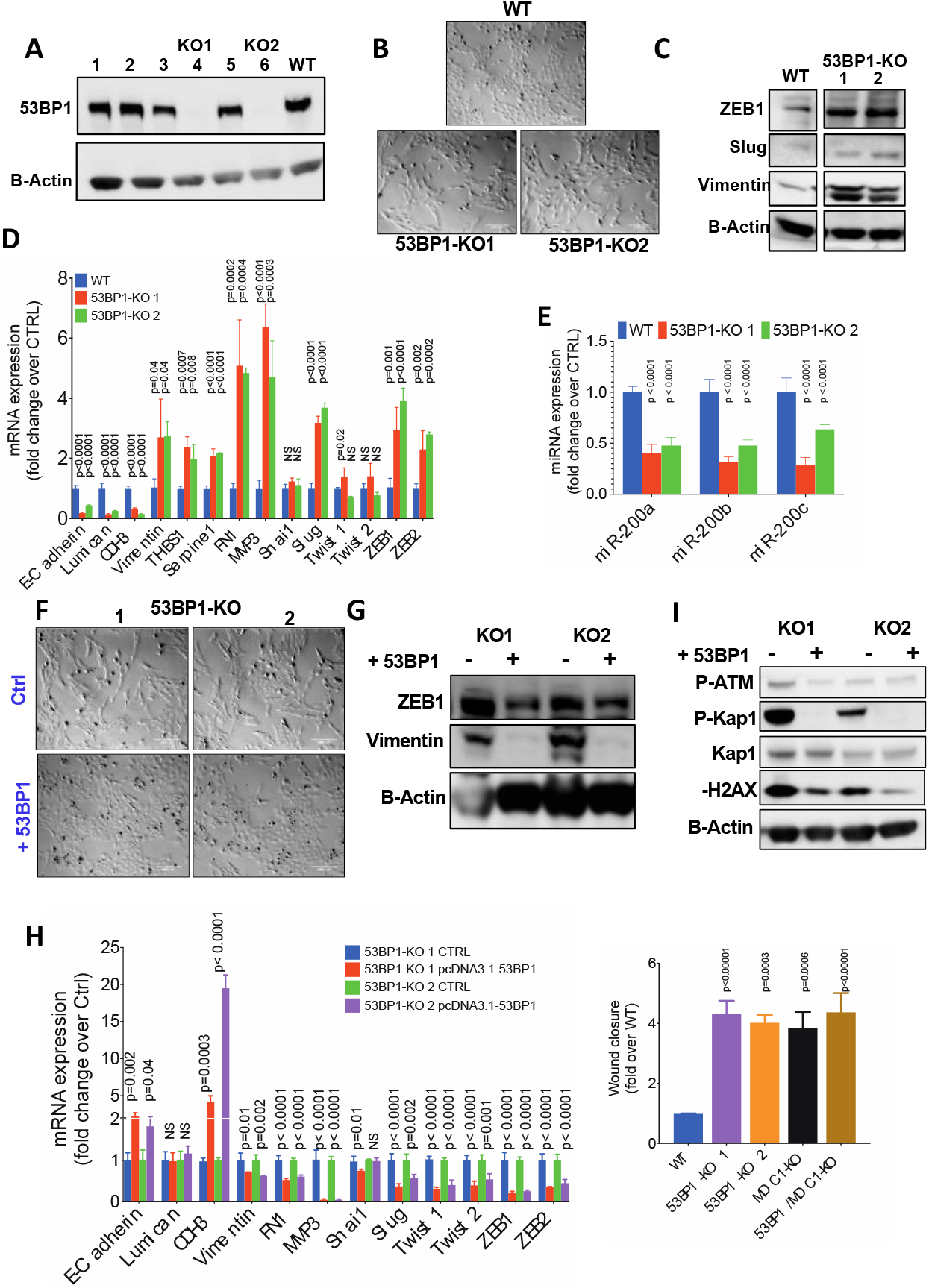
Disabling the DNA Damage Response induces EMT. **A**. Identification by western blot of CRISPR-Cas9 transfected HEK-Early clones with full KO for 53BP1 (KO1 and KO2). **B**. Cell morphology aspect of KO clones compared with the parental cell line (WT). Scale bar=200µm. **C**. Western blot analyses for EMT-related markers in 53BP1-KO clones. **D**. Relative mRNA level of expression for epithelial and EMT markers in 53BP1-KO clones compared to the parental cell line (WT). p values from comparisons with control are indicated. NS: non-significant. **E**. Relative expression level of three members of the miR-200 family upon deletion of 53BP1. p values from comparisons with control are indicated. **F**. Cell morphology aspect of 53BP1-KO clones before and after forced expression of exogenous 53BP1 (+53PB1). Scale bar=200µm. **G**. Western blot analyses showing the impact, on EMT-related markers, of 53BP1 re-expression in 53BP1-KO clones. **H**. Relative mRNA level of expression of epithelial and EMT markers in 53BP1-KO clones upon forced expression of exogenous 53BP1. p values from paired comparisons with cells transfected with a control (CTRL) plasmid are indicated. 000NS: non-significant. **I**. Western blot analyses showing the evolution of DNA damage markers β-ATM, β-KAP1 and β-H2AX in 53BP1-KO clones upon forced expression of exogenous 53BP1. **J**. Quantification of wound healing assays carried out with 53BP1-KO, MDC1-KO and 53BP1/MDC1-DKO cells. Represented are means of relative wound closures by KO clones after 20h in comparison to the parental cell line (WT). Original images are presented in Supplementary Figure S3G. p values correspond to pairwise comparisons with the parental cell line (WT).

We noticed that restoring the expression of 53BP1 in these cells was also associated to a decrease in the spontaneous levels of DNA damage markers P-H2AX, P-KAP1, and P-ATM (Figure 1I) suggesting that elimination of one component of the DDR in cells results in a paradoxical exacerbation of DDR, potentially explained by an accumulation of spontaneous DNA lesions. To confirm that abrogation of a DDR component spontaneously leads to EMT, we targeted MDC1, another DNA damage sensor that also accumulates at DSBs (*23*), for inactivation using the same Cas9 (D10A)-based approach. HEK-Early cells (53BP1 WT as well as 53BP1-KO cells) were transformed, sorted and cloned as above and one clone KO for MDC1 was identified in each context (Supplementary figure S2A). In agreement with results described for 53BP1-KO cells, MDC1-KO and MDC1/53BP1-double KO HEK cells displayed cell morphology changes compatible with EMT (Supplementary figure S2B) as well as enhanced expression of EMT-TFs and mesenchymal markers (Supplementary figure S2C-D), along with significant downregulation of both epithelial markers (Supplementary figure S2C) and EMT-related miRs (Supplementary figure S2E).

All DDR-KO HEK-Early clones were further characterized for proliferation and migration capacities, features expected to be affected by the EMT process. We detected a significant decrease in the proliferation rate of all KO clones in comparison with WT parental cells (Supplementary figure S2F), as previously shown for other cell lines undergoing EMT (*24*). Furthermore, all KO clones showed increased migration potential when compared to the parental cells, as evaluated in a wound healing assay (Figure 1J, Supplementary figure S2G).

Taken together, these results draw a strong link between DDR and EMT. They also support the notion that accumulation of endogenous damage (or exacerbation of DDR) – perhaps beyond a certain threshold-leads to EMT.

### EMT induction following 53BP1 abrogation occurs in a variety of carcinoma cells

We verified that EMT in response to the elimination of a DDR component was a general feature in epithelial tumor cells from different tissues. We knocked out 53BP1 in the epithelial colon cancer cell line HCT116 (either P53 WT or P53-KO), in the epithelial breast cancer cell line MCF7, and in a prostate epithelial cell line immortalized by SV40, PNT1A (Supplementary figure S3A). Deletion of 53BP1 in all these cell lines induced both loss of epithelial cell morphology (Supplementary figure S3B) and increase in gene expression of EMT-TFs and mesenchymal-related markers (Figure 2A-E), while epithelial markers and the miR-200 family all appeared to be downregulated (Figure 2A-F). Like HEK-Early cells, all cell lines displayed higher migration potential than their WT counterparts (Figure 2G) (Supplementary figure S3C). Together, these results confirm that spontaneous EMT because of DDR perturbation is a general phenomenon in transformed epithelial cells, and that this EMT may occur independently of the p53 status.

**Figure 2.**
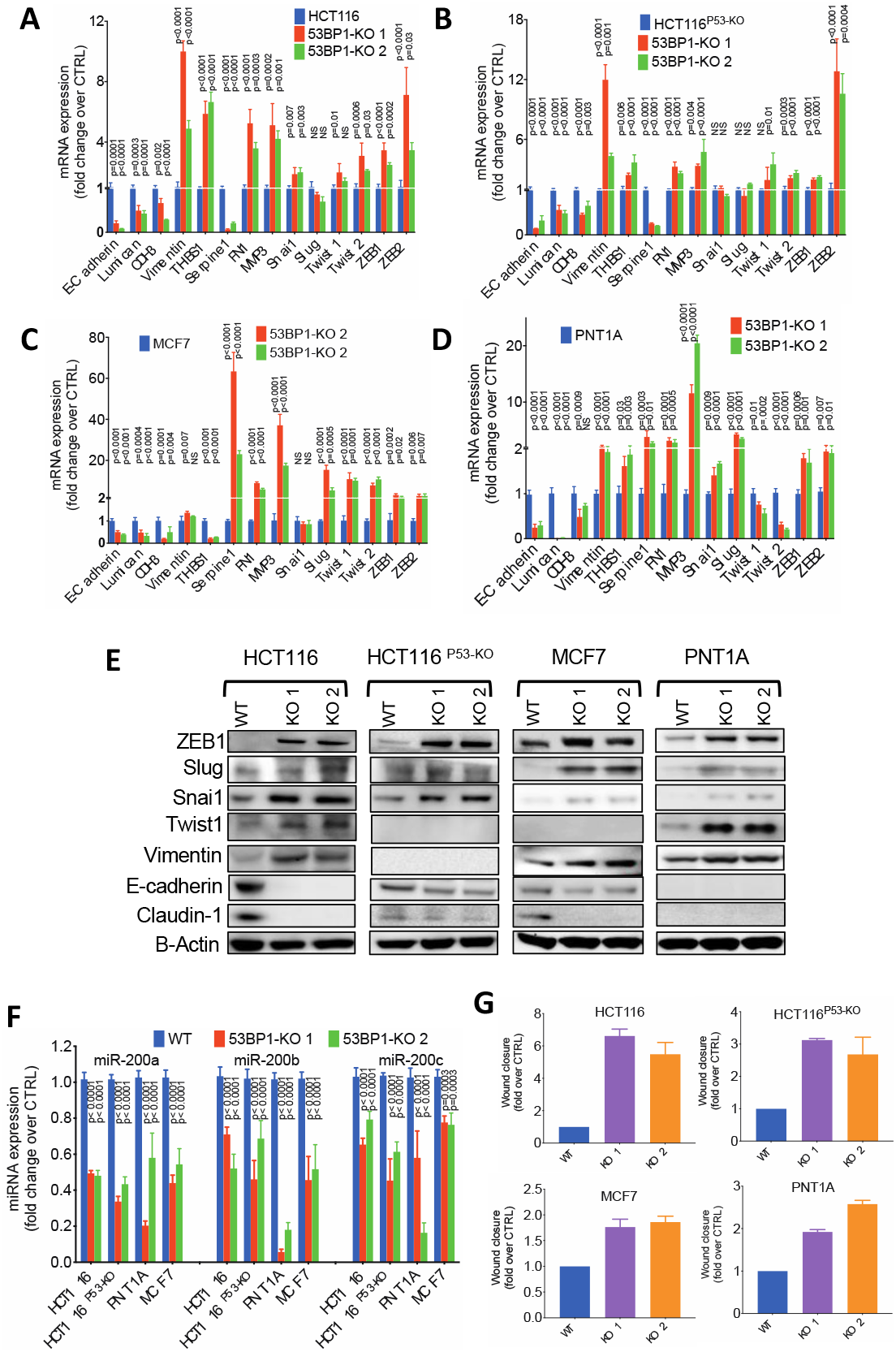
EMT induction following DDR attenuation is a widespread phenomenon among cancer cells. **A** to **D**. Relative mRNA level of expression of epithelial and EMT markers in 53BP1-KO clones isolated after transfection of four different cancer/transformed cell lines: HCT116 (colon, **A**), HCT116P53KO (same as HCT116 but KO for P53, **B**), MCF7 (breast, **C**) and PNT1A (prostate, **D**). In all cases, levels are compared to the parental cell line. p values from paired comparisons are indicated. NS: non-significant. **E**. Western blot analyses evaluating the level of expression of different EMT-related markers as well as epithelial markers in all four types of cells after deletion of 53BP1. **F**. Relative expression level of three members of the miR-200 family in all four types of cells after deletion of 53BP1. G. Quantification of wound healing assays carried out in all four cell types before and after deletion of 53BP1. Represented are relative wound closures in KO clones with respect to the parental cell line. Measurements were made after different times (indicated in Supplementary Figure **S3C** where original images are presented) depending on the parental cell’s wound healing capacity.

### Exposure to exogenous DNA damage reinforces EMT hallmarks in a DDR deficient context

The above results point to the possibility of a link between the levels of spontaneous DNA damage or DDR signaling found in a cell and the likelihood for that cell to undergo EMT. To determine whether further exposure to DNA damage could impact the DDR-related EMT phenotype already detected, we treated 53BP1-KO, and other knock-out cell clones, with etoposide 0.1 µM for 7 days and monitored the levels of EMT-related markers. As shown in Figure 3A (and Supplementary figure S4A), the mesenchymal aspect of KO cells was further reinforced. This phenomenon was coupled to a further increase in expression levels of EMT-TFs and other mesenchymal-related markers in 53BP1-KO (Figure 3B-C), as well as in MDC1-KO and MDC1/53BP1-KO (Supplementary figure S4B), while the expression of E-cadherin (albeit not that of Lumican) was further reduced. These results once again underscore the degree of plasticity of the EMT induced by the abrogation of one DDR component as well as the role played by DNA damage as a major trigger (or reinforcer) of EMT. They also formally establish that an intact DDR pathway is not absolutely required to escalate the EMT phenotypes in response to DNA damage.

**Figure 3.**
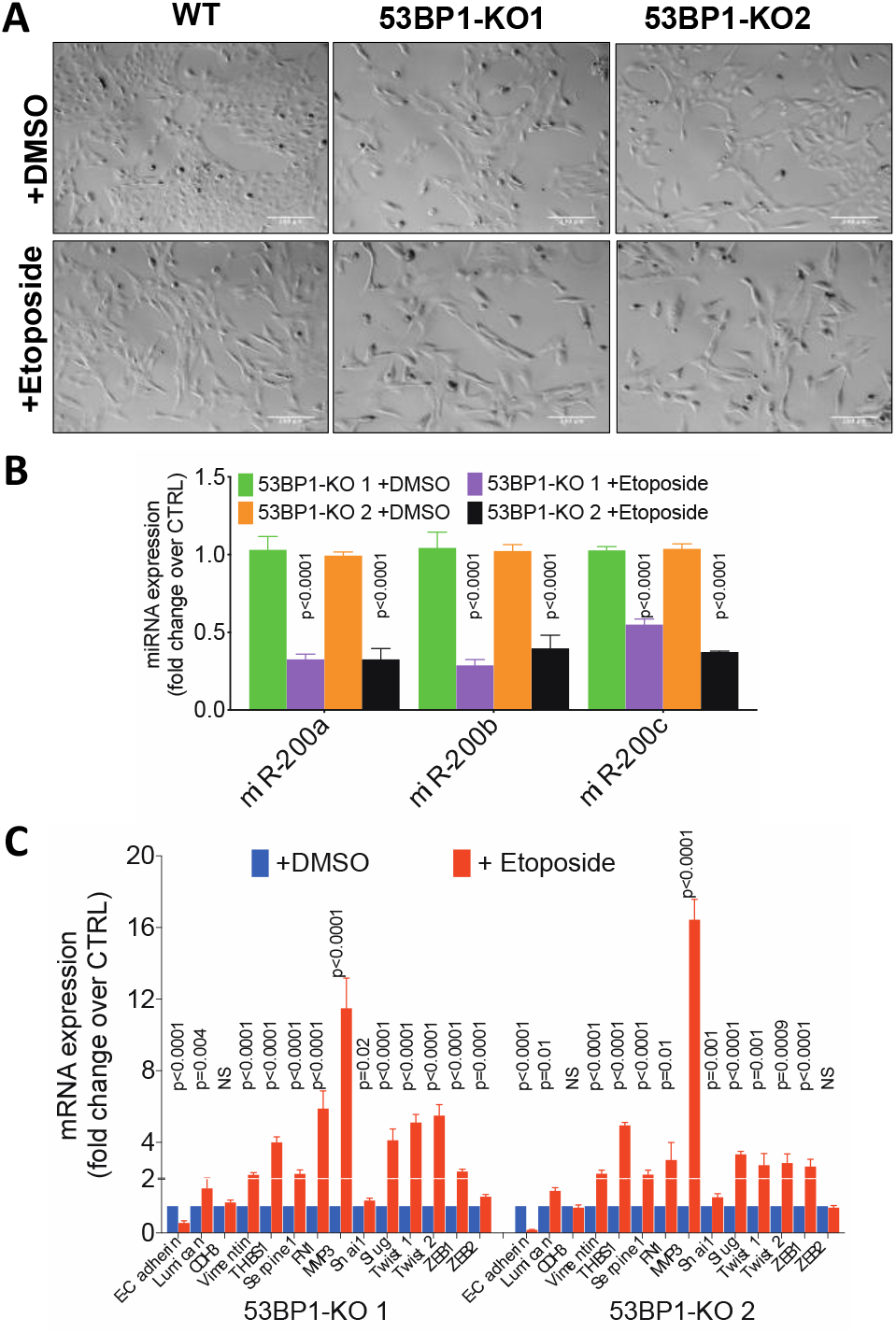
Exposure to exogenous DNA damage reinforces EMT hallmarks in a DDR deficient context. **A**. Cell morphology aspect of WT parental cell line and 53BP1-KO clones after treatment with etoposide 0.1µM for 7 days. Scale bar=200µm. **B**. Relative expression level of three members of the miR-200 family in 53BP1-KO clones after exposure to etoposide. p values from paired comparisons are indicated. **C**. Relative mRNA level of expression of epithelial and EMT markers in 53BP1-KO clones after exposure to etoposide. p values from paired comparisons are indicated. NS: non-significant.

### EMT promotes DNA repair

Our 53BP1 rescue experiment supported the notion that in cells partially disabled for DDR, the spontaneous exacerbation of DDR (likely due to accumulation of endogenous damage) triggers EMT. To further explore the relationship between DNA damage and EMT, we targeted the EMT-TFs ZEB1 for downregulation using RNA interference in both HCT116 and PNT1A 53BP1-KO cells. ZEB1 downregulation led, as expected, to downregulation in the expression of mesenchymal markers such as Vimentin and MMP3, and to upregulation of epithelial markers such as E-cadherin and Lumican in all KO cell lines (Figure 4A-B). Strikingly, the downregulation of ZEB1 also led to an increase in the levels of P-H2AX in HCT116 53BP1-KO (Figure 4C), a phenomenon that we also observed when Slug was targeted for downregulation in the same cells (Supplementary figure 5A). This intriguing observation strongly suggested a role for EMT-TFs in the protection against the spontaneous accumulation of endogenous DNA damage. Another possibility is that EMT-TFs’ activities somehow curb the intensity of the DDR. To confirm the potential role of EMT-TFs in DDR, we exogenously overexpressed TWIST1 in PNT1A cells, which display a relatively high level of spontaneous P-H2AX. As expected, TWIST1 overexpression induced an increase of mesenchymal markers, such as Vimentin and a decrease in epithelial markers such as E-Cadherin (Figure 4D). Remarkably, PNT1A cells overexpressing TWIST1 also displayed a strong decrease in the level of P-H2AX, supporting the conclusion that EMT reprogramming either improves DNA repair towards spontaneous DNA damage or restrain the levels of DDR (Figure 4D).

**Figure 4.**
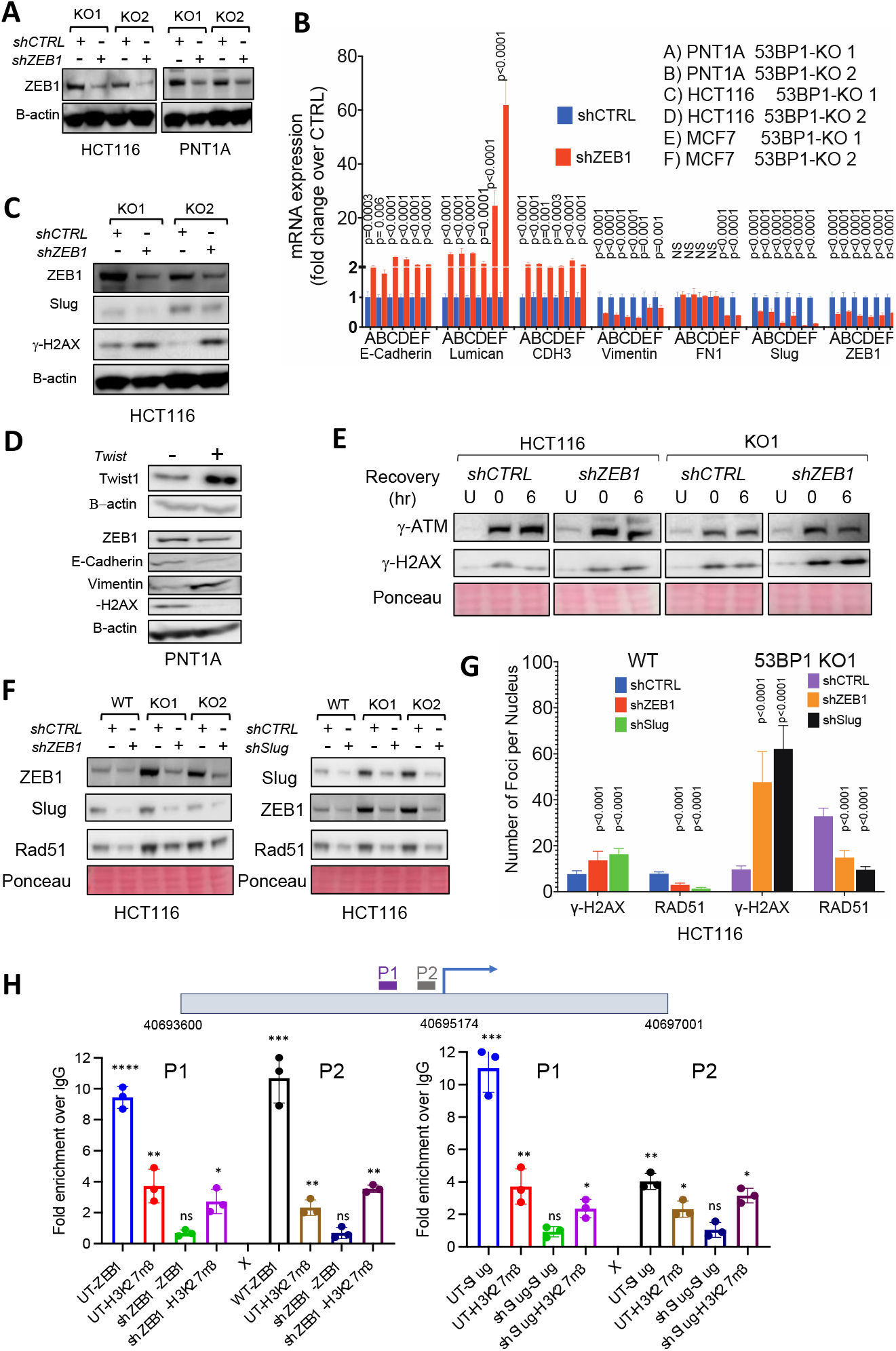
EMT favors DNA repair efficiency. **A**. Western blot analyses for ZEB1 after introduction of an shZEB1 expressing vector into different 53BP1-KO clones. **B**. Relative mRNA level of expression of epithelial and EMT markers in 53BP1-KO clones expressing shZEB1 as compared to the parental cell line (shCTRL). p values from paired comparisons are indicated. NS: non-significant. **C**. Western blot analyses showing the evolution of the DNA damage marker γH2AX in HCT116 53BP1-KO clones expressing shZEB1 as compared to the control (shCTRL). Also shown are the effects of this expression on ZEB1 and Slug. **D**. Western blot analyses showing the impact on EMT markers as well as the DNA damage marker γH2AX of forced expression of TWIST1 in PNT1A cells. **E**. DNA damage recovery assay in HCT116 cells, WT and KO for 53BP1, and depleted or not for ZEB1. Cells were treated (or not, U) with etoposide (2 µM for 24h) before medium change (0 hr) and let to recover for 6 hours. Shown here are western blot analyses for DNA damage markers γ-ATP and γH2AX. **F**. Western blot analyses for ZEB1, Slug and RAD51 after introduction of a vector expressing either shZEB1 or shSlug into WT or 53BP1-KO HCT116 cells. **G**. Quantification of γH2AX and RAD51 foci detected by immunofluorescence in WT or 53BP1-KO HCT116 cells expressing either shZEB1 or shSlug. p values from paired comparisons against the control (shCTRL) are indicated. **H**. Results from Cut&Run experiments using HCT116 cells and either anti-ZEB1 (bottom left) or anti-Slug (bottom right) antibodies for chromatin immunoprecipitation followed by enrichment analyses of two amplified DNA fragments (P1, P2) within the promoter region (ENSEMBL coordinates are indicated) of the RAD51 gene, shown here to span two small regions right upstream of the transcription initiation site (indicated by an arrow). HCT116 cells were either untreated (UT) or transduced with vectors expressing either shZEB1 (bottom left) or shSlug (bottom right). Positive control included an antibody against the histone mark H3K27me3. All PCR signals were normalized to the result obtained with positive antibody control (H3K27m3) and then to an amplified Alu fragment. Significant differences between WT and shZEB1 and shSLUG from multiple t-tests (adjusted) are indicated. ^*^, *p* < 0.05; ^**^, *p* < 0.01; ^***^, *p* < 0.001; ^****^, *p* < 0.0001.

To assess a potential role of EMT more directly in DNA repair, we dynamically evaluated the efficiency of this repair in HCT116 cells, WT or 53BP1-KO cells, depleted for ZEB1 and treated with etoposide for 24h. Cells were allowed to recover without the drug for 6 hours and the level of residual damage was determined by detecting P-H2AX. As shown in Figure 4E, P-H2AX levels decreased at 6h in control cells, while in both WT and 53BP1-KO ZEB1-depleted cells, the level of P-H2AX increased instead. We confirmed these observations in MCF7 cells, WT and 53BP1-KO, depleted or not for Slug (Supplementary figure S5B). In this case, cells were treated with etoposide and let to recover for 2 and 6 hours. Immunoblot analyses showed a rapid decrease in the levels of P-H2AX in control cells, while in cells (WT or 53BP1-KO) depleted for Slug, this decrease took longer (Supplementary figure S5B). These experiments demonstrate for the first time a link between EMT-TFs and DNA repair efficiency, including in tumor cells with intact DDR.

### EMT-TFs directly stimulate RAD51 transcription thus promoting DNA repair

To better characterize the relationship between EMT-TFs and DNA repair, we examined the consequences of EMT-TFs downregulation on the levels of RAD51, a key actor of HR-mediated repair, the other major repair pathway. Immunoblot analyses suggested that RAD51 levels are slightly increased in unperturbed 53BP1-KO cells with respect to parental cells (Figure 4F). Upon EMT-TF depletion, the levels of Rad51 decreased in both WT and 53BP1-KO HCT116 cells (Figure 4F), suggesting that these EMT-TFs control the expression of the Rad51 gene not only when cells undergo EMT but also in the epithelial state. The decrease on RAD51 expression upon knockdown of EMT-TFs and the concomitant increase in the levels of P-H2AX strongly argue in favor of an accumulation of spontaneous DNA damage in these cells as opposed to a damage-independent exacerbation of DDR (Figure 4C, Supplementary figure S5A). These results were confirmed in immunofluorescence experiments, where we detected a significant decrease of spontaneous RAD51 foci upon depletion of EMT-TFs factors, both in WT and 53BP1-KO HCT116 cells, even though this depletion was associated with a significant increase in P-H2AX foci (Figure 4G, Supplementary figure S5C). Finally, we examined the impact of EMT-TF depletion on drug resistance. While MCF7 53BP1-KO cells WT were more resistant to the drug than MCF7 WT (Supplementary figure S6A), both MCF7 53BP1-KO and WT cells depleted for ZEB1 or Slug were more sensitive than the control situation (Supplementary figure S6B-D). Together, these results indicate that the drug resistance characteristically associated with EMT may be, at least in part, explained by the impact of EMT-TFs on RAD51 expression leading to more efficient DNA repair by HR.

Thus far, our results point to a positive role for EMT in the repair of both endogenous and exogenous DNA damage and suggest that 53BP1-KO EMT cells, which are expected to be deficient in NHEJ, rather repair damage using HR. To draw a direct link between EMT and activation of HR, we explored the possibility that EMT-TFs themselves could bind to the RAD51 promoter. Cut-and-run experiments were performed in HCT116 cells, unperturbed or depleted for ZEB1, using anti-ZEB1 antibodies (and anti-H3K27me3 antibodies as a positive control) followed by q-PCR that targeted two different regions of the RAD51 promoter (Figure 4H), as well as the promoter of vimentin, a known target for EMT-TFs as a positive control (Supplementary Figure S6E). We detected a >5-fold enrichment signal for ZEB1 on the RAD51 promoter, with respect to no antibody control, and this enrichment was completely abrogated in cells partially depleted for ZEB1 (Figure 4H). Similar results were obtained using an anti-Slug antibody in the same cells, unperturbed or depleted for Slug (Figure 4H). As expected, we also obtained enrichments for the Vimentin promoter with both anti-ZEB1 and anti-Slug antibodies (Supplementary Figure S6E). These results demonstrate that EMT-TFs promote DNA repair through a direct stimulation of RAD51 transcription.

### Access to chromatin by EMT-TFs upon DNA damage depends on PARP and ALC1 activities

DNA repair requires increased DNA accessibility to repair factors and this accessibility depends on Poly (ADP-ribose) polymerase (PARP) activation and the subsequent activity of the PAR-binding chromatin remodeler CHD1L (ALC1)(*25, 26*). We wondered whether the same accessibility mechanisms were at play in the recruitment of EMT-TFs to promoters following DNA damage. To explore this possibility, we expressed a GFP-tagged ZEB1 version in U2OS cells, which were then subjected to DNA damage induced by laser microirradiation under the microscope. As shown in Figure 5A, the fusion protein rapidly accumulates at the sites of damage. Strikingly, treatment of the cells with olaparib, a PARP1/2 inhibitor in clinical use (*27*) and known to block chromatin remodeling by ALC1 upon DNA damage(*26*), prevents the rapid recruitment of ZEB1 to the sites of damage. Similar results were obtained in a cell context in which ALC1 had been abrogated (Figure 5B). These observations are compatible with the notion that chromatin relaxation mediated by ALC1 upon activation of PARP1 is a key step for EMT-TFs to access cognate sequences in response to DNA damage.

**Figure 5.**
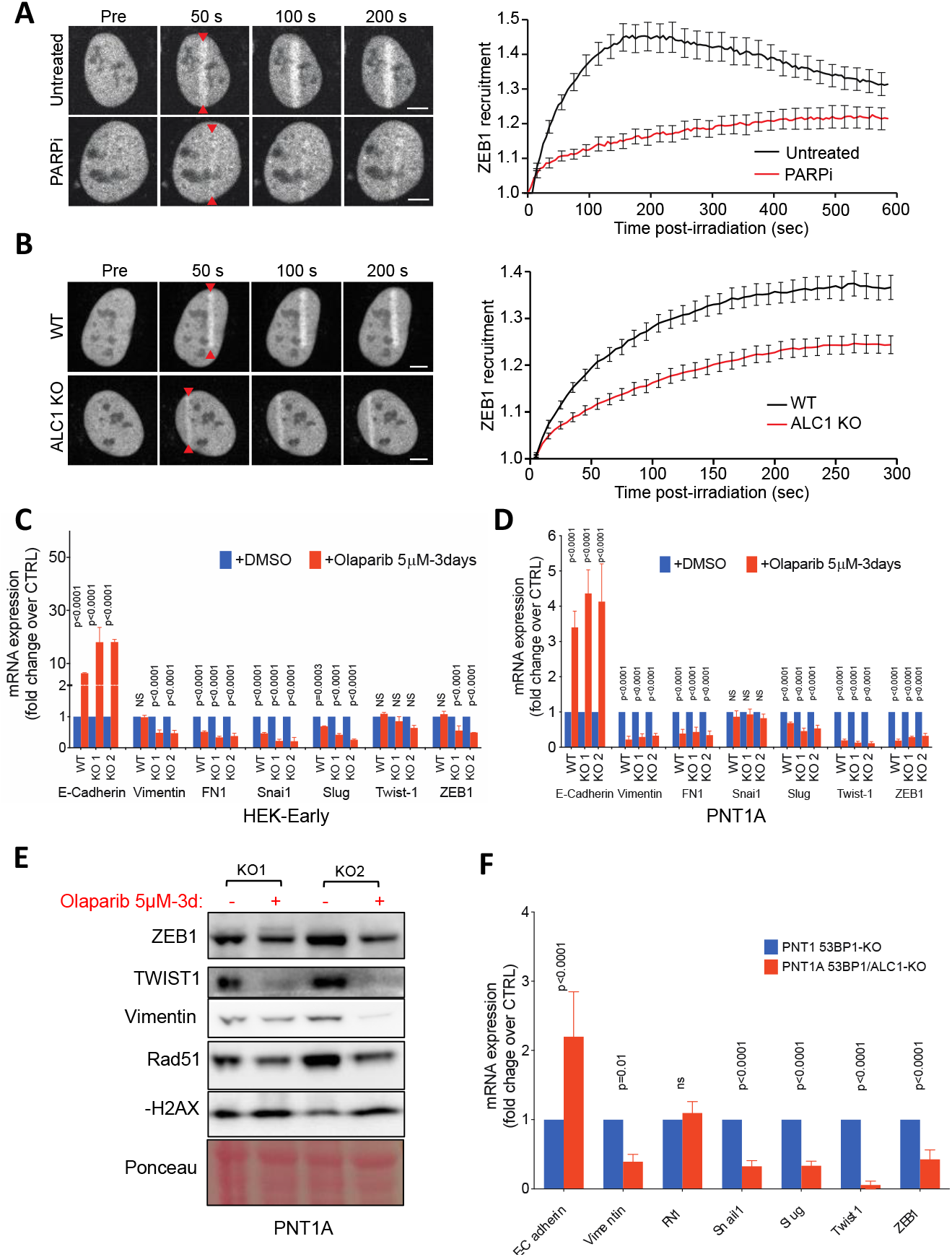
PARP/ALC1 are required for EMT-TF access to chromatin upon DNA damage. **A**. U2OS cells expressing a GFP-ZEB1 fusion protein were micro-irradiated with laser under the microscope across nuclei and the recruitment of the fluorescent protein was followed and quantified at different time points. Cells were treated or not with Olaparib, a PARP inhibitor, at 5µM. **B**. Similar experiments were carried out with U2OS cells that had been inactivated for ALC1. **C-D**. Relative mRNA level of expression of epithelial and EMT markers in 53BP1-KO HEK-Early cells (**C**) and in 53BP1-KO PNT1A cells (**D**) treated with Olaparib at the dose and time indicated, as compared to untreated cells. p values from paired comparisons are indicated. NS: non-significant. **E**. Western blot analyses of EMT-TFs, Vimentin, RAD51 and γH2AX in 53BP1-KO PNT1A cells before and after treatment with Olaparib. **F**. Relative mRNA level of expression of epithelial and EMT markers in 53BP1-KO PNT1A cells that were also inactivated for ALC1. p values from paired comparisons are indicated. NS: non-significant.

### Inhibition of PARP completely reverses EMT

We next tested the effect of olaparib on the EMT features presented by cells with DDR deficiency. Treatment with olaparib for only 3 days was sufficient to induce a strong reversal of EMT in HEK-early 53BP1-KO cells (Figure 5C). This rapid and complete reversal of EMT by olaparib was confirmed in PNT1A 53BP1-KO cells, which showed strong re-expression of E-cadherin and deep repression of both vimentin and fibronectin, as well as of most EMT-TFs (Figure 5D). Interestingly, olaparib treatment also induced a reinforcement in E-cadherin expression and a further decrease of mesenchymal markers (including EMT-TFs) in DDR-proficient epithelial transformed cells (Figures 5C-D). In agreement with our observations demonstrating a mechanistic link between EMT-TFs and RAD51 expression to stimulate repair, PNT1A 53BP1-KO cells treated with olaparib showed a decreased in the expression of RAD51 with an increase in -H2A accumulation, all in association with a decrease in ZEB1 and TWIST1 proteins (Figure 5E).

To verify that the DDR-related EMT also requires ALC1, we deleted the gene in PNT1A 53BP1-KO cells. As shown in Figure 5F, abrogation of ALC1 led to re-expression of E-cadherin and to suppression of major EMT-TFs and mesenchymal markers in these cells. These observations are consistent with the idea that PARP-dependent chromatin relaxation is required for EMT-TFs activity to induce EMT and stimulate RAD51 expression and that inhibitors of PARP1/2 could be very efficient in reversing EMT hallmarks associated with a defective DDR.

### Olaparib restores drug sensitivity to DDR-deficient EMT transformed cells

BRCA-deficient cancer cells are exquisitely sensitive to PARP inhibitors (*28*). We carried out cell survival tests for DDR-deficient EMT cells in the presence of olaparib which revealed that these cells were also more sensitive to the inhibitor than DDR-proficient cells (Supplementary figure S7A-B). Interestingly, although DDR-deficient cells were more resistant than WT cells to drugs like cis-platin or taxol (Figure 6A-D), in agreement with their EMT status, the combination of olaparib with these drugs restored their sensitivity (Figure 6A-D), also in agreement with the reversal of EMT hallmarks induced by olaparib. These experiments demonstrate that inhibition of PARP can overturn the resistance to chemotherapeutic agents usually associated with EMT.

**Figure 6.**
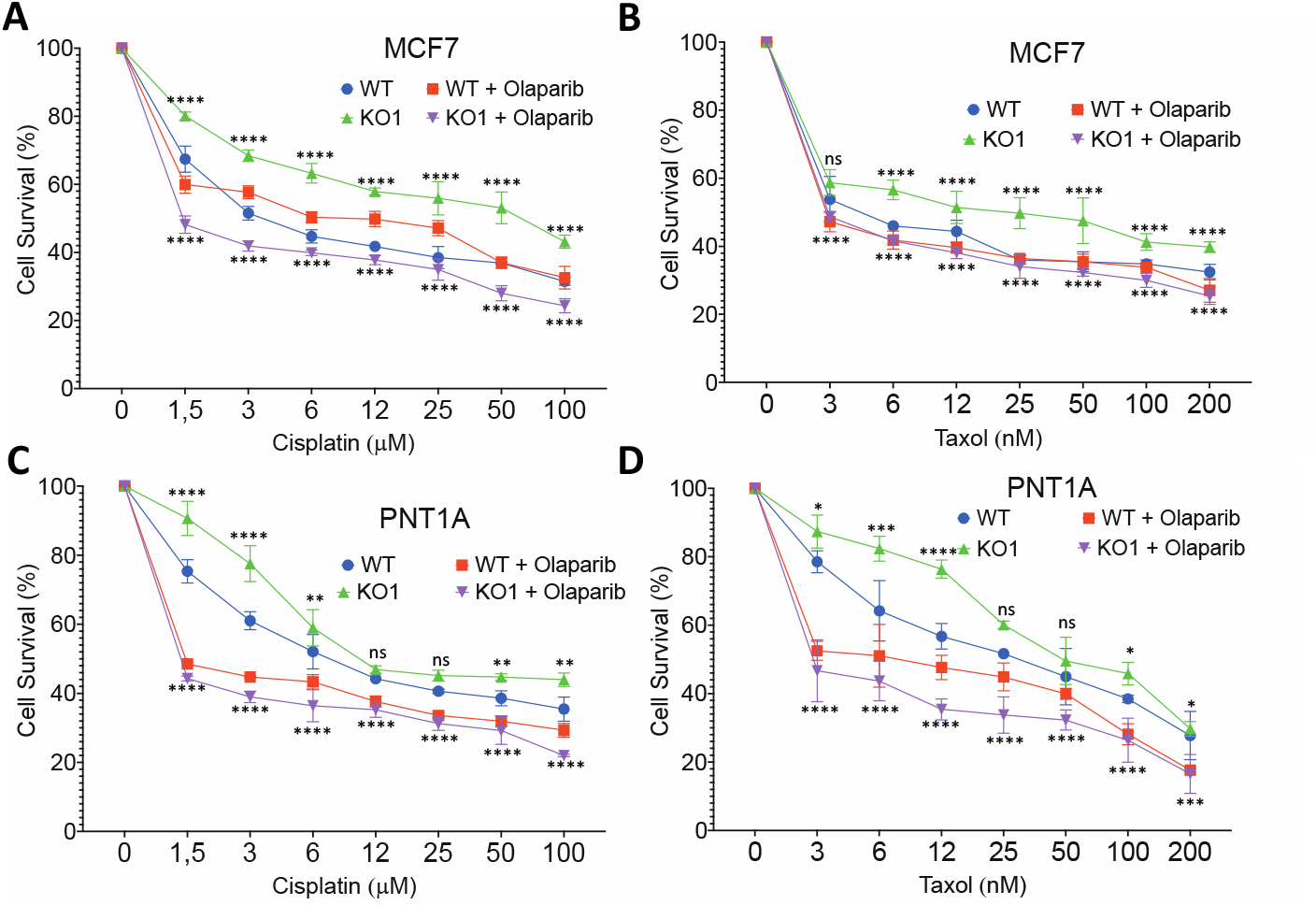
Olaparib restores drug sensitivity to DDR-deficient EMT transformed cells. **A to D**. Cytotoxicity assays for WT and 53BP1-KO MCF7 and for WT and 53BP1-KO PNT1A cells using cisplatin or taxol, alone or in combination with Olaparib (10 µM). Significant differences from multiple t-tests (adjusted) is indicated as follows: ^*^, *p* < 0.05; ^**^, *p* < 0.01; ^***^, *p* < 0.001; ^****^, *p* < 0.0001.

### EMT-TFs and RAD51 mRNA expression levels show positive correlation in most human cancers

To investigate whether the link between EMT-TFs and RAD51 had any potential clinical significance, we compared the levels of expression of mRNAs coding for EMT-TFs to those coding for RAD51 in cancerous and normal tissues, using publicly available RNAseq data from the TCGA consortium. The force of expression of transcripts coding for ZEB1, ZEB2, SNAI1, SNAI2 (Slug), TWIST1 and TWIST2, was considered together and was correlated to the force of expression detected for RAD51 in the same sample (http://gepia2.cancer-pku.cn/#index). Compellingly, 32 out of 34 paired correlations, which comprised 18 different types of cancers, showed a statistically significant positive correlation, with many of them being highly significant (Table 1). This remarkable result highlights the mechanistic relationship between EMT and DNA repair in clinically relevant tumor contexts.

**Table 1.** Expression levels of EMT transcriptional factors are positively correlated to the expression levels of RAD51 in most human cancers. RNAseq expression data was obtained from publicly available TCGA data. Comparisons were made using a Spearman’s correlation. P-Value and R-value are shown

| TCGA Cancer |  |  |  | TCGA Normal |  |  |
| --- | --- | --- | --- | --- | --- | --- |
| Cancer name | EMT Gene | P-Value | R | EMT Gene | P-Value | R |
| ACC (adrenocortical carcinoma) | Slug | 9.2e-05 | 0.43 | Slug | 0.035 | 0.19 |
| ACC (adrenocortical carcinoma) | ZEB1 | 0.0035 | 0.33 | ZEB1 | 0.14 | 0.13 |
| ACC (adrenocortical carcinoma) | ZEB2 | 0.00036 | 0.4 | ZEB2 | 0.81 | -0.021 |
| BLCA (Bladder urothelial carcinoma) | Snail1 | 0.00065 | 0.17 | Snail1 | 0.35 | -0.23 |
| ESCA (Esophageal carcinoma) | Twist1 | 0.014 | 0.18 | Twist1 | 0.22 | -0.36 |
| ESCA (Esophageal carcinoma) | Slug | 0.00094 | 0.24 | Slug | 0.46 | -0.23 |
| GBM (Glioblastoma multiforme) | ZEB1 | 2.8e-13 | 0.53 | ZEB1 | 0.25 | 0.11 |
| GBM (Glioblastoma multiforme) | ZEB2 | 0.002 | 0.24 | ZEB2 | 0.6 | 0.051 |
| LGG (lower grade glioblastoma) | Twist1 | 1.3e-13 | 0.32 | ZEB1 | 0.25 | 0.11 |
| LGG (lower grade glioblastoma) | ZEB1 | 3.7e-10 | 0.27 | ZEB2 | 0.93 | 0.008 |
| LIHC (liver hepatocellular carcinoma) | ZEB1 | 5.3e-06 | 0.23 | ZEB1 | 0.91 | 0.017 |
| LUAD (Lung adenocarcinoma) | Snail1 | 4.1e-14 | 0.33 | Snail1 | 0.49 | -0.093 |
| LUAD (Lung adenocarcinoma) | Twist1 | 6.3e-12 | 0.31 | Twist1 | 0.99 | 0.001 |
| MESCO (mesothelioma) | Slug | 6.7e-05 | 0.41 |  |  |  |
| MESCO (mesothelioma) | Twist1 | 0.0031 | 0.31 |  |  |  |
| MESCO (mesothelioma) | ZEB2 | 0.036 | 0.22 |  |  |  |
| OV (Ovary tumor) | Snail1 | 9.7e-07 | 0.23 | Snail1 | 0.71 | -0.041 |
| PAAD (Prostate adenocarcinoma) | Twist1 | 3.0e-04 | 0.27 | Twist1 | 0.53 | -0.47 |
| PAAD (Prostate adenocarcinoma) | Slug | 7.1e-05 | 0.29 | Slug | 0.37 | -0.63 |
| READ (Rectum adenocarcinoma) | Slug | 0.085 | 0.18 | Slug | 0.13 | -0.51 |
| READ (Rectum adenocarcinoma) | ZEB2 | 0.1 | 0.17 | ZEB2 | 0.014 | -0.74 |
| SKCM (Skin cutaneous melanoma) | ZEB2 | 5.3e-06 | 0.21 | ZEB2 | 0.38 | -0.058 |
| STAD (stomach adenocarcinoma) | Snail1 | 0.0028 | 0.15 | Snail1 | 0.02 | -0.39 |
| TGCT (testicular germ cell tumors) | Snail1 | 9.9e-12 | 0.54 | Snail1 | 3.3e-07 | -0.38 |
| THCA (Thyroid carcinoma) | Slug | 1.3e-08 | 0.25 | Slug | 0.41 | -0.11 |
| THCA (Thyroid carcinoma) | ZEB2 | 4.1e-09 | 0.26 | ZEB2 | 0.018 | 0.14 |
| THYM (Thyoma) | ZEB1 | 6.2e-08 | 0.47 | ZEB1 | 1 | -1 |
| UCS (Uterine carcinosarcoma) | Slug | 0.022 | 0.3 | Slug | 0.33 | 0.11 |
| UCS (Uterine carcinosarcoma) | Snail1 | 0.0061 | 0.36 | Snail1 | 0.93 | -0.011 |
| UCS (Uterine carcinosarcoma) | ZEB1 | 0.0067 | 0.36 | ZEB1 | 0.81 | -0.027 |
| UVM (Uveal melanoma) | ZEB1 | 4.0e-06 | 0.49 | ZEB1 | 1.5e-07 | -0.29 |
| UVM (Uveal melanoma) | ZEB2 | 0.0073 | 0.3 | ZEB2 | 0.0036 | -0.16 |
| UVM (Uveal melanoma) | Twist2 | 0.022 | 0.26 | Twist2 | 0.012 | -0.14 |
| UVM (Uveal melanoma) | Snail1 | 0.043 | 0.23 | Snail1 | 0.93 | -0.011 |

## Discussion

Here, we have provided further evidence that transformed epithelial cells exposed to DNA damage undergo EMT. More importantly, we have demonstrated that this response is entirely dependent on PARP/ALC1 activation, which then allows EMT-TFs to access cognate promoters to initiate reprogramming. These results provide for the first time a mechanistic link between DNA damage and EMT. Unexpectedly, we also show in this work that RAD51, a key component of DNA repair by homologous recombination, is a major target of EMT-TFs thus allowing us to propose that stimulation of homologous recombination is a key aspect of the drug-resistance seen in tumor epithelial cells that undergo EMT.

Intriguingly, disabling 53BP1 or MDC1, key sensors that initiate DDR, and as such expected to participate in the EMT response, was spontaneously associated with an enduring EMT, which also depended on PARP/ALC1 activation. Our observations strongly suggest that the trigger of this reprogramming when DDR is abrogated appears to be the persistence of endogenous, unrepaired, DNA damage as suggested by the spontaneous increase in the levels of phosphorylation of DNA damage sensors such as ATM, KAP1 and H2AX. These results agree with a published work showing that abolishing H2AX, another DNA damage sensor, promotes EMT in HCT116 cells (*20*). Strikingly, 53BP1-KO, MDC1-KO and 53BP1/MDC1-DKO cells were still able to respond to additional exogenous DNA damage by reinforcing EMT-related traits, which could also be prevented by inhibiting PARP/ALC1 activation, thus underlying the extent of the plasticity acquired by cells under genomic stress.

How do DDR-deficient cells sense persistent DNA damage? Our experiments demonstrate that the PARP enzymatic activity is tightly involved in the sensing and transduction of the signal that leads to EMT. The role of PARP in EMT is not unprecedented since work by other groups already have pointed out that PARP inhibitors are able to prevent TGF-β-induced EMT in tumors cells (*29-31*), thus supporting the contention that PARP acts downstream from TGF-β activation. Given the role of PARP1 both as a DNA damage sensor and in the choice of DNA repair pathway, PARP activity appears to be the key component of the DDR-related EMT signaling pathway. Furthermore, inhibiting PARP1/2 or abrogating ALC1 were quite efficient in reversing already established DDR-related EMT phenotypes. Interestingly, this EMT reversion also encompasses a reversal of the enhanced resistance initially conferred by the EMT to clinically relevant chemotherapeutic drugs such as cis-platin and taxol. Finally, transgenic animal models have suggested that, opposite to what has been described when PARP activity is inhibited, obliteration of the PARP1 gene leads to increased TGF-β and EMT during prostate tumorigenesis (*31*). Potential explanations to this apparent conundrum include the fact that, although PARP1 is responsible for most of the ADP-ribosylation detected in cells, ADP-ribosylation in response to genotoxic stress is not totally abolished in PARP1-KO cells (*32*). In fact, the presence of significant amounts of ADP-ribosylation in mouse embryonic fibroblasts from a PARP1-KO strain led to the discovery of PARP2 (*33*), the other nuclear PARP enzyme whose activity in also enhanced in response to DNA damage (*34*). Indeed, simultaneous and specific targeting of both PARP1 and PARP2 by olaparib (*35*) would explain why this inhibitor very efficiently prevents/reverses DNA damage-related EMT phenotypes.

More generally, our results offer a sound mechanistic explanation helping to understand how EMT is triggered during tumorigenesis. Indeed, DNA replication stress is omnipresent in cancer cells (*36, 37*), which could provide a persistent source of endogenous DNA damage and as shown in this work, a trigger to trans-differentiation. Such replication stress is expected to occur early during tumorigenesis as oncogene activation leads to premature entry into S-phase and to activation of ectopic replication origins, thus creating replication/transcription conflicts ultimately leading to fork collapse and double strand breaks (*38*). Indeed, signs of DNA damage response is often detected in precancerous lesions, followed by the detection of chromosome instability in early stages of tumor development (*38*). Our data clearly point to the possibility that DNA damage related EMT may occur as soon as the tumorigenic process initiates and that tumor cells may acquire metastatic properties at early stages of cancer progression. This hypothesis is supported by the demonstration of disseminated cancer cells (DCCs) in patients with breast cancer before any metastasis is detected (*39*) and in transgenic mouse models of breast and pancreatic cancers as well as melanoma (*40-42*).

The present work also holds potential clinical implications. EMT has been previously directly connected to drug resistance (*43*). Our data suggest that exposure to chemotherapeutic drugs could, on its own, induce an EMT-like state in tumor cells in vivo. Importantly, also as shown here, this DNA damage-related EMT response not only can be prevented but also reversed if PARP1/2 activity is inhibited. Of note, PARP1/2 inhibitors like olaparib have been approved for the treatment of BRCA mutated ovarian and breast cancers (*27*) because HR-deficient tumor cells display an exquisite sensitivity to these inhibitors, which block the other major repair pathway NHEJ (*44*). This has led to the idea that cancer patients with mutations in other genes that participate in HR repair (such as ATM, ATR, BAP1, CDK12, CHEK2, FANCA, FANCC, FANCD2, FANCE, FANCF, PALB2, NBS1, WRN, RAD51C, RAD51D, MRE11A, CHEK1, BLM, and RAD51B) could benefit from association treatments using PARP1/2 inhibitors (*45*). Here, we show that tumor cells deficient for 53BP1 (a major signaling molecule for NHEJ) also display increased sensitivity to PARP1/2 inhibitors. This apparently counterintuitive result may be explained by the fact that, as shown experimentally here and further supported by the significant co-expression of EMT-TFs and RAD51 in a variety of human cancers, EMT promotes DNA repair by stimulating RAD51 expression and the HR pathway. Therefore, PARP inhibitors can affect both major DNA repair pathways, particularly in cells that have undergone an EMT.

Altogether, our work provides experimental as well as theoretical grounds to envision enlargements of current studies on the potential applications of PARP inhibitors in cancer treatment schemes. For instance, these studies may explore the benefits of an association of PARP inhibitors as early as possible in a majority of, if not all, carcinoma types or at least whenever DDR is intrinsically affected or is the target of a therapeutic intervention.

## Materials and Methods

### Cell lines and cell culture

HA5-early cells obtained from HEK (Human Epithelial Kidney) cells, as described by Castro-Vega et al.,(*5*). The PNT1A cell line was derived from human post-pubertal prostate normal epithelial cells transformed with SV40 (*46*). HCT116 parental cells and HCT116 P53-KO cells were obtained as gift from Dr. Antonin Morillon and Dr. Franck Toledo, respectively. MCF7 and U2OS cells were obtained from ATCC. U2OS ALC1-KO cells were described previously(*26*). All cell lines were cultured at 37 °C in a humidified atmosphere containing 5% CO2. HA5-early, PNT1A, and MCF7 parental and transfection derived cell lines were cultured in Minimum Essential Medium-alpha 1X (MEM-α 1X) (Gibco; Life technologies), supplemented with 10% (v/v) fetal bovine serum (FBS; Eurobio), 1% (v/v) MEM Non-essential Amino Acid Solution 100X (Gibco; Life technologies), and 1% (v/v) sodium pyruvate 100X (Gibco; Life technologies). HCT116 parental and transfected cells were cultured in McCoy’s (5A)-modified (Gibco; Life technologies) supplemented with 10% (v/v) fetal bovine serum. U2OS cells were cultured in DMEM (4.5 g/l glucose, Sigma) supplemented with 10% fetal bovine serum (Life Technologies), 2 mM glutamine (Sigma), 100 µg/ml penicillin and 100 U/ml streptomycin (Sigma).

### All-in-one nickase plasmids

All-in-One-GFP and All-in-One-mCherry plasmids were purchased from Addgene (AIO-GFP #74119 and AIO-mCherry #74120). Their constructions have been described in (*22*).

### Golden Gate assembly of CRISPR sgRNA cloning

53BP1 and MDC1 sgRNAs were inserted in All-in-one nickase plasmids backbone as described by Chiang et al.(*22*) and the insertion and presence of both sgRNA sequences were verified by DNA Sanger sequencing in a single reaction using a primer: 5′-CTTGATGTACTGCCAAGTGGGC-3′. sgRNA sequences for 53BP1 and MDC1 are shown in Supplementary Table 1.

### Transfection and transduction

To obtain H2AX-KO, 53BP1-KO, MDC1-KO, double 53BP1/MDC1-KO HA5-early cells, and 53BP1-KO PNT1A and MCF7 cells, parental cells were transfected with All-in-One nickase plasmids via electroporation using the amaxa-nucleofector® II according to the manufacturer’s instructions (Thermofisher). 5-10 μg of the All-in-One plasmid were transfected into 10^6^ cells. Cell Line NucleofectorTM Kit V (Catalog #: VCA-1003) (Lonza) was used for transfection procedure according to the manufacturer’s instructions. Q-001 program was applied to transfect. To obtain 53BP1-KO, HCT116 (including WT and P53-KO) cells were transfected using jetPRIME® (Reference number: 114-07) (Polyplus) according to the manufacturer’s instructions. To obtain a knockdown of EMT-TFs in cells, HCT116, PNT1A, MCF7 parental and 53BP1-KO cells were transfected with shZEB1 and shSlug (a kind gift from Pr. Alain Puisieux lab) and control shRNAs using jetPrime according to manufacturer’s instructions. CRISPR knockouts were all verified by genotype screening and immunoblot analysis. shRNA Knockdown transfections followed by at least 3 weeks of selection by puromycin treatment. To obtain 53BP1 restored expressing cells, HA5-Early WT and 53BP1-KO cells were transduced with lentiviral particles packaged in 293T cells using the plasmid 53BP1.pcDNA3.1 (kind gift from Dr. Gaëlle Legube, University of Toulouse). Then selection of transduced cells continued for three weeks under treatment with Hygromycin 75 µg/ml concentration. To obtain TWIST1 overexpressing clone from PNT1A cell line, cells were transfected with Twist (TWIST1) (NM_000474) Human Tagged ORF Clone (Origene). Transfected cells underwent 3 weeks selection procedure with Neomycin (400 µg/ml). For transient expression of GFP-tagged ZEB1, U2OS cells were transfected 12–24 h after seeding into eight-well Imaging Chamber CG (Zell-Kontakt) with XtremeGENE HP (Sigma) according to the manufacturer’s instructions and incubated for 48 h prior to imaging.

### Fluorescence-activated cell sorting (FACS)

72 hours after transfection, cells were trypsinized, washed with PBS, resuspended in PBS (supplemented with 2% FBS) and filtered through cell strained capped 5 ml falcon (STEM CELL). Cells were individually sorted, either based on GFP, mCherry, or mCherry+EGFP markers into TPP tissue culture testplate-96 (SIGMA-ALDRICH) at a single-cell-per-well density for clonal expansion. Subsequently, all viable clones passed through the genotyping screening and Immunoblot analysis.

### CRISPR KO clone selection

Single cell sorting following each transfection was continued by propagation of clones in P96-well plate and passage into P24-well plate, then into P6-well plate. Cells pellets were collected after washing and trypsinization for DNA and protein analysis. Clones were verified by immunoblotting after initial genotype screening and DNA sequencing indicating rearrangement of the relevant locus. When possible, at least two clones were randomly selected for this study.

### Genomic DNA extraction

For genotype screening, genomic DNA was extracted from clones in a 96-well plate format. To extract DNA, cells were washed with 200 μl PBS and incubated in 50 μl Lysis buffer (10mM Tris pH 7.5, 10mM EDTA, 10 mM NaCl, 0.5 % sarcosyl) for overnight at 55°C. DNA precipitated and attached to the surface by adding 100 µl chilled EtOH + 75mM NaCl and incubated for at least 30 minutes at room temperature. The lysates were removed by traversing the plate. DNA was washed by incubation with 150 µl 70% EtOH. EtOH was removed and DNA after solved in 30 µl DEPC‐treated water. For DNA sequencing, genomic DNA was extracted using the Qiamp DNA mini kit (QIAGEN) according to the manufacturer’s instructions.

### PCR and Genotype screening

All PCRs in this study were performed with Promega PCR master kit according to the manufacturer’s instructions. PCR products were separated on the agarose gels and analyzed by Geldoc XR+ system (Biorad). PCR products were sequenced by using gene-specific primers. A list of the PCR primers used for genotypic screening or DNA sequencing is shown in Supplementary Table S2.

### Immunoblotting

Proteins were extracted with lysis buffer (1% Triton, and 50 mM Tris-HCl, pH 8.0, 300 µM NaCl, 5 mM EDTA). Protein concentrations were determined with a BMG Labtech FluoStar Optima Plate Reader at 560 nm. Samples were then heated to 95 °C for 5 minutes following the addition of loading dye. Proteins separated by NuPAGE™ 4-12% Bis-Tris precast protein gels (Thermo Fisher Scientific), and then transferred to nitrocellulose membrane (GE Healthcare). To confirm homogeneous loading and transferring, membranes were stained with Ponceau S. Then Ponceau S stained membranes were washed with PBS tween followed by 1-hour incubation with blocking solution 5% BSA-PBS tween or 5% milk-PBS tween. Then the membrane was incubated overnight with primary antibodies diluted in blocking solution. The primary antibody was washed with PBS tween and the membrane was incubated with specific secondary antibody diluted in blocking solution. Antibodies used in this study are listed in Supplementary Table S3.

### Quantitative RT-PCR

RNeasy mini kit (Qiagen) was used to extract RNA from cells, according to the manufacturer’s instructions. RNA was resuspended in DEPC‐treated water. Synthesis of cDNA with Superscript III reverse transcriptase (Invitrogen) was primed with oligo (dT).

Primer sequences are shown in Supplementary Table S4. Analyses were carried out using Promega PCR master mix (Promega) on an LC-480 Real-Time PCR system (Roche). Amounts of target mRNA were normalized to an endogenous housekeeping gene (γ-actin) and calculated as fold change over control or parental cell lines.

To perform QRT-PCR analysis for miRNAs, miRNeasy kit was used to extract total RNA and microRNAs. Then miRCURY LNA™ Universal RT microRNA cDNA Synthesis Kit (ThernoFiscer) used for RT PCR according to the manufacturer instructions. miRCURY LNA SYBR® Green PCR Kit (Qiagen) was used for QRT-PCR according to manufacturer instructions and as described in (*5*).

### Proliferation assays

Cells were plated in cell culture P12 plate at a density of 10,000 cells in triplicates. Every 24 hours wells were trypsinized and resuspended in media. 200µl of cell suspension was diluted in 10ml of isotone solution and counted by Beckman Coulter Z2 cell/particle counter. The last counting is performed 96 hours after cell plating.

### Wound healing or migration assays

Cells were plated and grown to full confluence in 6-well plates. Pipette tips were used to create scratches in the monolayers. Pictures were taken by an inverted light microscope in T0 (beginning of the experiment) and T1, variable time depending on the cell line (between 6 and 24 hours) after wounding.

### Cell survival assays

10,000 Cells per well were plated and grown for 24 hours in 96-well plates. Cells were treated in serial diluted concentrations of each drug indicated in the text. Treatments lasted 96 hours. After 4 days, density of cells in each well was quantified using Methylene Blue staining. In the first step, cells (wells) were washed with PBS 1X. Then 100 µl absolute methanol is added to each well and plate was incubated for 1 hour at room temperature. Then the wells were let to dry and 100 µl methylene blue solution (concentration 1gr/L) was added to each well and followed by 1-hour incubation at room temperature. Following the staining step, wells were rinsed with water 2 times and then let the wells to dry. Washing step followed by solubilization by adding 200 µl HCL (0.1N) in each well and incubation at 60°C for 30 minutes. In the last step O.D of each well was measured at 630 nm using BMG FLUOstar OPTIMA plate reader.

### Drug treatments

For each treatment experiment, cells were plated at an appropriate density 24 hours prior to treatment. The dosages indicated in the text for each treatment were selected according to the cell lines specific IC50 values and duration of the treatment period. Each drug treatment was refreshed every 48-72 hours according to the manufacturer indications. Treatment was removed and followed by washing with PBS 1X to collect the pellet to perform QRT-PCR and/or Immunoblotting. The following drugs were used in this study: Etoposide (S1225) (Selleckchem), and Olaparib (AZD2281) (Selleckchem), Cisplatine (Mylan, 100mg/100ml) and Taxol (Paclitaxel) (S1150) (Selleckchem).

### Etoposide treatment recovery assays

100K Cells per well were plated and grown to for 24 hours in 6-well plates. Cells were treated with the indicated Etoposide concentrations and incubated for the mentioned time. Following the removal of Etoposide treatment, wells were washed with PBS 1X and collected by trypsinization after the indicated time following the removal of treatment.

### IF cell-staining and microscopy-based screening

Cells were washed with PBS containing 0.1% Tween-20 (PBST) and fixed for 15 min in PBS containing 2% paraformaldehyde. After fixation, cells were permeabilized in PBS containing 0.2–0.5% Triton X-100 for 10–15min and blocked with blocking buffer (PBST with 5% (w/v) BSA) for 30min. After three washes with PBST, immunostaining was performed with the indicated primary antibodies (Supplementary Table S3) diluted in the blocking buffer for 1 hour at room temperature followed by three washes with PBST. Cells were then stained with the appropriate secondary antibodies for 1h at room temperature diluted in blocking buffer followed by another three washes with PBST. Cells were counterstained with DAPI (1 μg/ml) and mounted using Vectashield (Vector Labs). Images were acquired with a Zeiss Axio Imager Fluorescence Microscope. Images were analyzed using Image J.

### Cleavage under targets and release using nuclease (CUT&RUN)

CUT&RUN experiment according to the protocol described by Skene el al(*47*). ZEB1 (Origene, # TA802298) and Slug (Origene, # OTI1A6) antibodies were used. As a positive control α-H3K27me3 rabbit monoclonal antibody (Cell Signaling Technology, # 9733). As a negative control, mouse IgG antibody was used. DNA precipitation followed by DNA purification according to the same protocol. QPCR was done for Rad51 promoter using three sets of primers (see Supplementary Table 4) and data normalized to Alu as reference and then made ratio to positive control.

### Fluorescence imaging, laser microirradiation and ZEB1 recruitment kinetics to DNA lesions

Prior to live cell imaging combined with laser irradiation, U2OS cells were presensitized with 0.3 µg/ml Hoechst 33342 for 1 h at 37°C. Then, growth medium was replaced with CO_2_-independent imaging medium (Phenol Red-free Leibovitz’s L-15 medium (Life Technologies) supplemented with 20% fetal bovine serum, 100 µg/ml penicillin and 100 U/ml streptomycin). For PARP inhibition, cells were treated with 30 µM Olaparib (Euromedex) for 30 min prior to imaging. Live-cell imaging experiments combined with laser irradiation were performed on a Ti-E inverted microscope (Nikon) equipped with a CSU-X1 spinning-disk head (Yokogawa). The cells were imaged with a Plan APO 60×/1.4 NA oil-immersion objective lens and a sCMOS ORCA Flash 4.0 camera. The fluorescence of EGFP was excited at 490 and detected with a bandpass filter at 500-550 nm. Laser microirradiation was performed along a 16 µm-line through the nucleus using a single-point scanning head (iLas2 from Roper Scientific) coupled to the epifluorescence backboard of the microscope. To ensure reproducibility, laser power at 405 nm was measured at the beginning of each experiment and set to 125 µW at the sample level. Cells were maintained at 37°C with a heating chamber in absence of CO2. These experiments were completed with unsynchronized cells. The recruitment kinetics were quantified with FIJI (https://fiji.sc/). The images sequences were first registered using the MultiStackReg plugin(*48*), then regions of interest were delineated manually to measure the mean fluorescence intensity in the damaged region (*I*_*d*_), in the whole nucleus (*I*_*n*_) and in a background region outside of the cell (*I*_*bg*_). Protein accumulation at sites of damage (*A*_*d*_) was then calculated as:

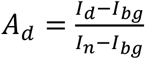

The intensity within the microirradiated area was then normalized to the intensity prior to damage induction.

### Statistical analysis

Data were obtained from a minimum of 3 independent experiments. For PCR, each experiment was done in triplicate. In bar graphs and dose-response curves, comparisons between control and test samples were performed using either two-way Anova test, Student’s *t* test or multiple *t* tests. All statistical tests used a two-tailed α = 0.05 level of significance and were performed using GraphPad Prism (GraphPad Software). For all studies, curves show pooled data with error bars representing SEM obtained from at least three independent experiments. For cytotoxic curves, significance is indicated in the graphs as follows: ^*^, *P* < 0.05; ^**^, *P* < 0.01; ^***^, *P* < 0.001; ^****^, *P* < 0.0001.

## Funding

Work in the Londono-Vallejo laboratory has been supported by grants from:

The Fondation ARC (ALV).

The French Institut National du Cancer (INCa) (ALV)

The INCa-Ligue-ARC PAIR prostate cancer program (ALV). Institut Curie post-doctoral program (FR).

## Author contributions

FR has planned and carried out most of experiments. W-YLB, MS and RS carried out specific experiments. SH and AL-V conceived and planned experiments. FR, RS, SH and AL-V analyzed the data, prepared the figures and tables and wrote the paper. All authors have read and approved the manuscript.

## Competing interests

Authors declare that they have no competing interests.

## Data and materials availability

All data (original pictures or excel files) and materials used in the analyses are available. All data are available in the main text or the supplementary materials.

## Notes

### Competing Interest Statement

The authors have declared no competing interest.

## References

1. R. Derynck, R. A. Weinberg, EMT and Cancer: More Than Meets the Eye. Dev Cell 49, 313–316 (2019).

2. M. P. Stemmler, R. L. Eccles, S. Brabletz, T. Brabletz, Non-redundant functions of EMT transcription factors. Nat Cell Biol 21, 102–112 (2019).

3. A. Puisieux, T. Brabletz, J. Caramel, Oncogenic roles of EMT-inducing transcription factors. Nat Cell Biol 16, 488–494 (2014).

4. I. Pastushenko, C. Blanpain, EMT Transition States during Tumor Progression and Metastasis. Trends Cell Biol 29, 212–226 (2019).

5. L. J. Castro-Vega, K. Jouravleva, W. Y. Liu, C. Martinez, P. Gestraud, P. Hupe, N. Servant, B. Albaud, D. Gentien, S. Gad, S. Richard, S. Bacchetti, A. Londono-Vallejo, Telomere crisis in kidney epithelial cells promotes the acquisition of a microRNA signature retrieved in aggressive renal cell carcinomas. Carcinogenesis 34, 1173–1180 (2013).

6. L. J. Castro-Vega, K. Jouravleva, P. Ortiz-Montero, W. Y. Liu, J. L. Galeano, M. Romero, T. Popova, S. Bacchetti, J. P. Vernot, A. Londono-Vallejo, The senescent microenvironment promotes the emergence of heterogeneous cancer stem-like cells. Carcinogenesis 36, 1180–1192 (2015).

7. H. Rajagopalan, C. Lengauer, CIN-ful cancers. Cancer Chemother Pharmacol 54 Suppl 1, S65–68 (2004).

8. R. S. Maser, R. A. DePinho, Connecting chromosomes, crisis, and cancer. Science 297, 565–569 (2002).

9. I. M. Shih, W. Zhou, S. N. Goodman, C. Lengauer, K. W. Kinzler, B. Vogelstein, Evidence that genetic instability occurs at an early stage of colorectal tumorigenesis. Cancer Res 61, 818–822 (2001).

10. S. F. Bakhoum, B. Ngo, A. M. Laughney, J. A. Cavallo, C. J. Murphy, P. Ly, P. Shah, R. K. Sriram, T. B. K. Watkins, N. K. Taunk, M. Duran, C. Pauli, C. Shaw, K. Chadalavada, V. K. Rajasekhar, G. Genovese, S. Venkatesan, N. J. Birkbak, N. McGranahan, M. Lundquist, Q. LaPlant, J. H. Healey, O. Elemento, C. H. Chung, N. Y. Lee, M. Imielenski, G. Nanjangud, D. Pe’er, D. W. Cleveland, S. N. Powell, J. Lammerding, C. Swanton, L. C. Cantley, Chromosomal instability drives metastasis through a cytosolic DNA response. Nature 553, 467–472 (2018).

11. M. A. Nowak, N. L. Komarova, A. Sengupta, P. V. Jallepalli, M. Shih Ie, B. Vogelstein, C. Lengauer, The role of chromosomal instability in tumor initiation. Proc Natl Acad Sci U S A 99, 16226–16231 (2002).

12. E. M. Kass, M. E. Moynahan, M. Jasin, When Genome Maintenance Goes Badly Awry. Mol Cell 62, 777–787 (2016).

13. S. Wolters, B. Schumacher, Genome maintenance and transcription integrity in aging and disease. Front Genet 4, 19 (2013).

14. H. T. Lynch, T. G. Shaw, J. F. Lynch, Inherited predisposition to cancer: a historical overview. Am J Med Genet C Semin Med Genet 129C, 5–22 (2004).

15. N. Romero-Laorden, E. Castro, Inherited mutations in DNA repair genes and cancer risk. Curr Probl Cancer 41, 251–264 (2017).

16. T. Watanabe, T. Kobunai, Y. Yamamoto, K. Matsuda, S. Ishihara, K. Nozawa, H. Yamada, T. Hayama, E. Inoue, J. Tamura, H. Iinuma, T. Akiyoshi, T. Muto, Chromosomal instability (CIN) phenotype, CIN high or CIN low, predicts survival for colorectal cancer. J Clin Oncol 30, 2256–2264 (2012).

17. B. S. Taylor, N. Schultz, H. Hieronymus, A. Gopalan, Y. Xiao, B. S. Carver, V. K. Arora, P. Kaushik, E. Cerami, B. Reva, Y. Antipin, N. Mitsiades, T. Landers, I. Dolgalev, J. E. Major, M. Wilson, N. D. Socci, A. E. Lash, A. Heguy, J. A. Eastham, H. I. Scher, V. E. Reuter, P. T. Scardino, C. Sander, C. L. Sawyers, W. L. Gerald, Integrative genomic profiling of human prostate cancer. Cancer Cell 18, 11–22 (2010).

18. E. I. Harper, E. F. Sheedy, M. S. Stack, With Great Age Comes Great Metastatic Ability: Ovarian Cancer and the Appeal of the Aging Peritoneal Microenvironment. Cancers (Basel) 10, (2018).

19. P. Ortiz-Montero, A. Londono-Vallejo, J. P. Vernot, Senescence-associated IL-6 and IL-8 cytokines induce a self- and cross-reinforced senescence/inflammatory milieu strengthening tumorigenic capabilities in the MCF-7 breast cancer cell line. Cell Commun Signal 15, 17 (2017).

20. U. Weyemi, C. E. Redon, R. Choudhuri, T. Aziz, D. Maeda, M. Boufraqech, P. R. Parekh, T. K. Sethi, M. Kasoji, N. Abrams, A. Merchant, V. N. Rajapakse, W. M. Bonner, The histone variant H2A.X is a regulator of the epithelial-mesenchymal transition. Nat Commun 7, 10711 (2016).

21. Y. Xu, D. Xu, Repair pathway choice for double-strand breaks. Essays Biochem, (2020).

22. T. W. Chiang, C. le Sage, D. Larrieu, M. Demir, S. P. Jackson, CRISPR-Cas9(D10A) nickase-based genotypic and phenotypic screening to enhance genome editing. Sci Rep 6, 24356 (2016).

23. Z. Lou, K. Minter-Dykhouse, X. Wu, J. Chen, MDC1 is coupled to activated CHK2 in mammalian DNA damage response pathways. Nature 421, 957–961 (2003).

24. H. J. Hugo, N. Gunasinghe, B. G. Hollier, T. Tanaka, T. Blick, A. Toh, P. Hill, C. Gilles, M. Waltham, E. W. Thompson, Epithelial requirement for in vitro proliferation and xenograft growth and metastasis of MDA-MB-468 human breast cancer cells: oncogenic rather than tumor-suppressive role of E-cadherin. Breast Cancer Res 19, 86 (2017).

25. R. Smith, T. Lebeaupin, S. Juhasz, C. Chapuis, O. D’Augustin, S. Dutertre, P. Burkovics, C. Biertumpfel, G. Timinszky, S. Huet, Poly(ADP-ribose)-dependent chromatin unfolding facilitates the association of DNA-binding proteins with DNA at sites of damage. Nucleic Acids Res 47, 11250–11267 (2019).

26. H. Sellou, T. Lebeaupin, C. Chapuis, R. Smith, A. Hegele, H. R. Singh, M. Kozlowski, S. Bultmann, A. G. Ladurner, G. Timinszky, S. Huet, The poly(ADP-ribose)-dependent chromatin remodeler Alc1 induces local chromatin relaxation upon DNA damage. Mol Biol Cell 27, 3791–3799 (2016).

27. J. Ledermann, P. Harter, C. Gourley, M. Friedlander, I. Vergote, G. Rustin, C. L. Scott, W. Meier, R. Shapira-Frommer, T. Safra, D. Matei, A. Fielding, S. Spencer, B. Dougherty, M. Orr, D. Hodgson, J. C. Barrett, U. Matulonis, Olaparib maintenance therapy in patients with platinum-sensitive relapsed serous ovarian cancer: a preplanned retrospective analysis of outcomes by BRCA status in a randomised phase 2 trial. Lancet Oncol 15, 852–861 (2014).

28. B. Evers, R. Drost, E. Schut, M. de Bruin, E. van der Burg, P. W. Derksen, H. Holstege, X. Liu, E. van Drunen, H. B. Beverloo, G. C. Smith, N. M. Martin, A. Lau, M. J. O’Connor, J. Jonkers, Selective inhibition of BRCA2-deficient mammary tumor cell growth by AZD2281 and cisplatin. Clin Cancer Res 14, 3916–3925 (2008).

29. M. Schacke, J. Kumar, N. Colwell, K. Hermanson, G. A. Folle, S. Nechaev, A. Dhasarathy, L. Lafon-Hughes, PARP-1/2 Inhibitor Olaparib Prevents or Partially Reverts EMT Induced by TGF-beta in NMuMG Cells. Int J Mol Sci 20, (2019).

30. O. Karicheva, J. M. Rodriguez-Vargas, N. Wadier, K. Martin-Hernandez, R. Vauchelles, N. Magroun, A. Tissier, V. Schreiber, F. Dantzer, PARP3 controls TGFbeta and ROS driven epithelial-to-mesenchymal transition and stemness by stimulating a TG2-Snail-E-cadherin axis. Oncotarget 7, 64109–64123 (2016).

31. H. Pu, C. Horbinski, P. J. Hensley, E. A. Matuszak, T. Atkinson, N. Kyprianou, PARP-1 regulates epithelial-mesenchymal transition (EMT) in prostate tumorigenesis. Carcinogenesis 35, 2592–2601 (2014).

32. L. Rank, S. Veith, E. C. Gwosch, J. Demgenski, M. Ganz, M. C. Jongmans, C. Vogel, A. Fischbach, S. Buerger, J. M. Fischer, T. Zubel, A. Stier, C. Renner, M. Schmalz, S. Beneke, M. Groettrup, R. P. Kuiper, A. Burkle, E. Ferrando-May, A. Mangerich, Analyzing structurefunction relationships of artificial and cancer-associated PARP1 variants by reconstituting TALEN-generated HeLa PARP1 knock-out cells. Nucleic Acids Res 44, 10386–10405 (2016).

33. J. C. Ame, V. Rolli, V. Schreiber, C. Niedergang, F. Apiou, P. Decker, S. Muller, T. Hoger, J. Menissier-de Murcia, G. de Murcia, PARP-2, A novel mammalian DNA damage-dependent poly(ADP-ribose) polymerase. J Biol Chem 274, 17860–17868 (1999).

34. Q. Chen, M. A. Kassab, F. Dantzer, X. Yu, PARP2 mediates branched poly ADP-ribosylation in response to DNA damage. Nat Commun 9, 3233 (2018).

35. C. E. Knezevic, G. Wright, L. L. R. Rix, W. Kim, B. M. Kuenzi, Y. Luo, J. M. Watters, J. M. Koomen, E. B. Haura, A. N. Monteiro, C. Radu, H. R. Lawrence, U. Rix, Proteome-wide Profiling of Clinical PARP Inhibitors Reveals Compound-Specific Secondary Targets. Cell Chem Biol 23, 1490–1503 (2016).

36. T. D. Halazonetis, V. G. Gorgoulis, J. Bartek, An oncogene-induced DNA damage model for cancer development. Science 319, 1352–1355 (2008).

37. S. K. Sotiriou, T. D. Halazonetis, Remodeling Collapsed DNA Replication Forks for Cancer Development. Cancer Res 79, 1297–1298 (2019).

38. V. G. Gorgoulis, L. V. Vassiliou, P. Karakaidos, P. Zacharatos, A. Kotsinas, T. Liloglou, M. Venere, R. A. Ditullio, Jr., N. G. Kastrinakis, B. Levy, D. Kletsas, A. Yoneta, M. Herlyn, C. Kittas, T. D. Halazonetis, Activation of the DNA damage checkpoint and genomic instability in human precancerous lesions. Nature 434, 907–913 (2005).

39. H. Hosseini, M. M. S. Obradovic, M. Hoffmann, K. L. Harper, M. S. Sosa, M. Werner-Klein, L. K. Nanduri, C. Werno, C. Ehrl, M. Maneck, N. Patwary, G. Haunschild, M. Guzvic, C. Reimelt, M. Grauvogl, N. Eichner, F. Weber, A. D. Hartkopf, F. A. Taran, S. Y. Brucker, T. Fehm, B. Rack, S. Buchholz, R. Spang, G. Meister, J. A. Aguirre-Ghiso, C. A. Klein, Early dissemination seeds metastasis in breast cancer. Nature 540, 552–558 (2016).

40. Y. Husemann, J. B. Geigl, F. Schubert, P. Musiani, M. Meyer, E. Burghart, G. Forni, R. Eils, T. Fehm, G. Riethmuller, C. A. Klein, Systemic spread is an early step in breast cancer. Cancer Cell 13, 58–68 (2008).

41. A. D. Rhim, E. T. Mirek, N. M. Aiello, A. Maitra, J. M. Bailey, F. McAllister, M. Reichert, G. L. Beatty, A. K. Rustgi, R. H. Vonderheide, S. D. Leach, B. Z. Stanger, EMT and dissemination precede pancreatic tumor formation. Cell 148, 349–361 (2012).

42. J. Eyles, A. L. Puaux, X. Wang, B. Toh, C. Prakash, M. Hong, T. G. Tan, L. Zheng, L. C. Ong, Y. Jin, M. Kato, A. Prevost-Blondel, P. Chow, H. Yang, J. P. Abastado, Tumor cells disseminate early, but immunosurveillance limits metastatic outgrowth, in a mouse model of melanoma. J Clin Invest 120, 2030–2039 (2010).

43. T. Shibue, R. A. Weinberg, EMT, CSCs, and drug resistance: the mechanistic link and clinical implications. Nat Rev Clin Oncol 14, 611–629 (2017).

44. H. E. Bryant, N. Schultz, H. D. Thomas, K. M. Parker, D. Flower, E. Lopez, S. Kyle, M. Meuth, N. J. Curtin, T. Helleday, Specific killing of BRCA2-deficient tumours with inhibitors of poly(ADP-ribose) polymerase. Nature 434, 913–917 (2005).

45. S. Boussios, C. Abson, M. Moschetta, E. Rassy, A. Karathanasi, T. Bhat, F. Ghumman, M. Sheriff, N. Pavlidis, Poly (ADP-Ribose) Polymerase Inhibitors: Talazoparib in Ovarian Cancer and Beyond. Drugs R D 20, 55–73 (2020).

46. A. Degeorges, F. Hoffschir, O. Cussenot, C. Gauville, A. Le Duc, B. Dutrillaux, F. Calvo, Recurrent cytogenetic alterations of prostate carcinoma and amplification of c-myc or epidermal growth factor receptor in subclones of immortalized PNT1 human prostate epithelial cell line. Int J Cancer 62, 724–731 (1995).

47. P. J. Skene, J. G. Henikoff, S. Henikoff, Targeted in situ genome-wide profiling with high efficiency for low cell numbers. Nat Protoc 13, 1006–1019 (2018).

48. P. Thevenaz, U. E. Ruttimann, M. Unser, A pyramid approach to subpixel registration based on intensity. IEEE Trans Image Process 7, 27–41 (1998).

