## Supplemental files for "DNA damage-induced PARP/ALC1 activation leads to Epithelial-to-Mesenchymal transition stimulating homologous recombination"

Supplementary Materials for

### PARP/ALC1 activation following DNA damage provokes Epithelial-to-Mesenchymal transition which fosters homologous recombination.

Fatemeh Rajabi et al

**This PDF file includes:**

Figs. S1 to S7
Tables S1 to S4

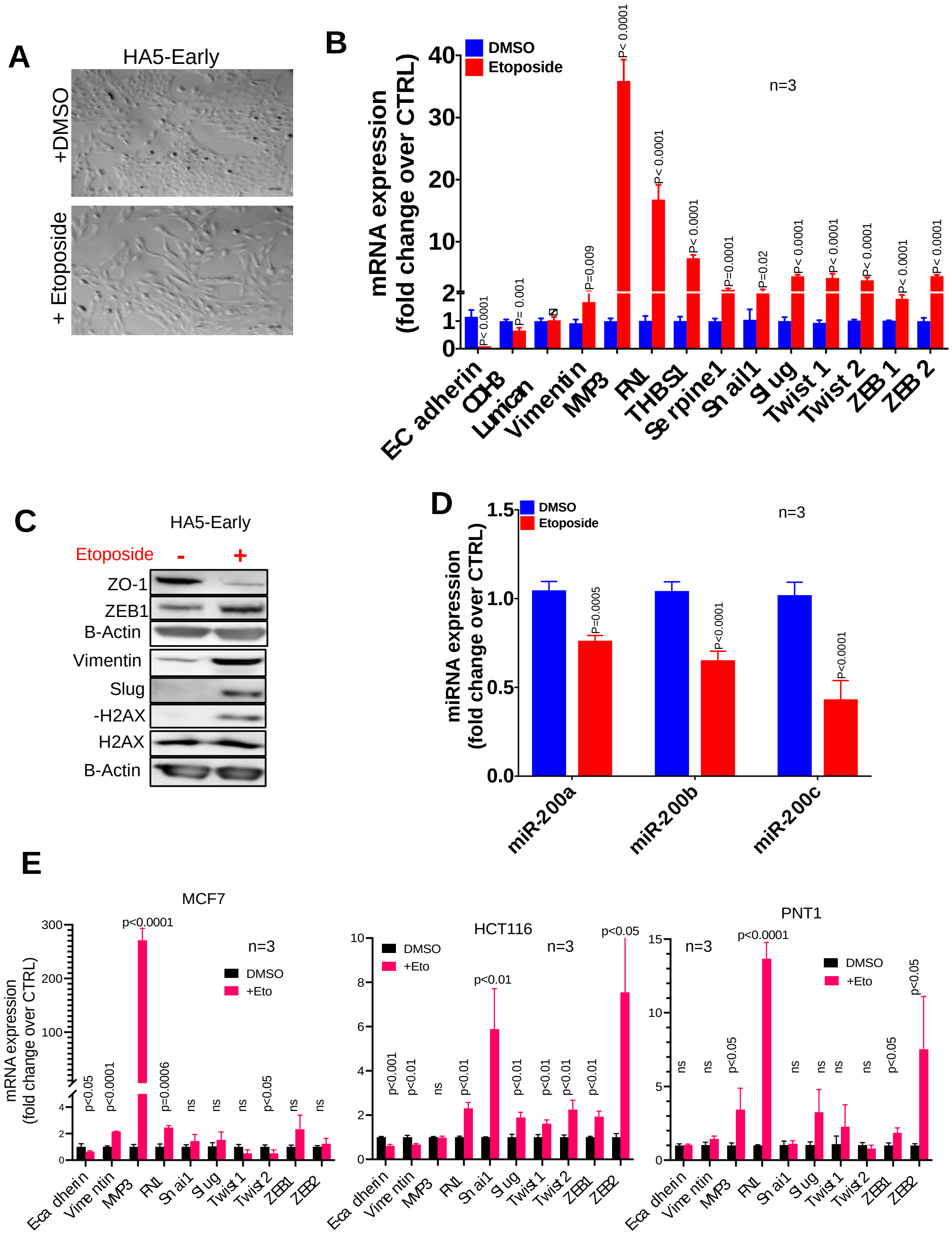


**Fig. S1.**

DNA damage rapidly induces EMT in transformed epithelial cells. **A.** Cell morphology of HEK cells before (DMSO) and after 7 days treatment with etoposide. **B.** Relative mRNA level of expression of epithelial and EMT markers in HEK cells treated with etoposide and compared to the untreated cell line (DMSO). p values from paired comparisons are indicated. ns: non-significant. **C**. Detection by western blot of some epithelial and mesenchymal markers in HEK cells before and after treatment with etoposide. **D.** Relative expression level of three members of the miR-200 family after treatment with etoposide and compared to untreated cells (DMSO). p values from paired comparisons are indicated. **E.** Relative mRNA level of expression of epithelial and EMT markers in three different tumor-derived (MCF7 and HCT116) or transformed (PNT1A) cells treated with etoposide and compared to the untreated counterparts (DMSO). p values from paired comparisons are indicated. ns: non-significant.

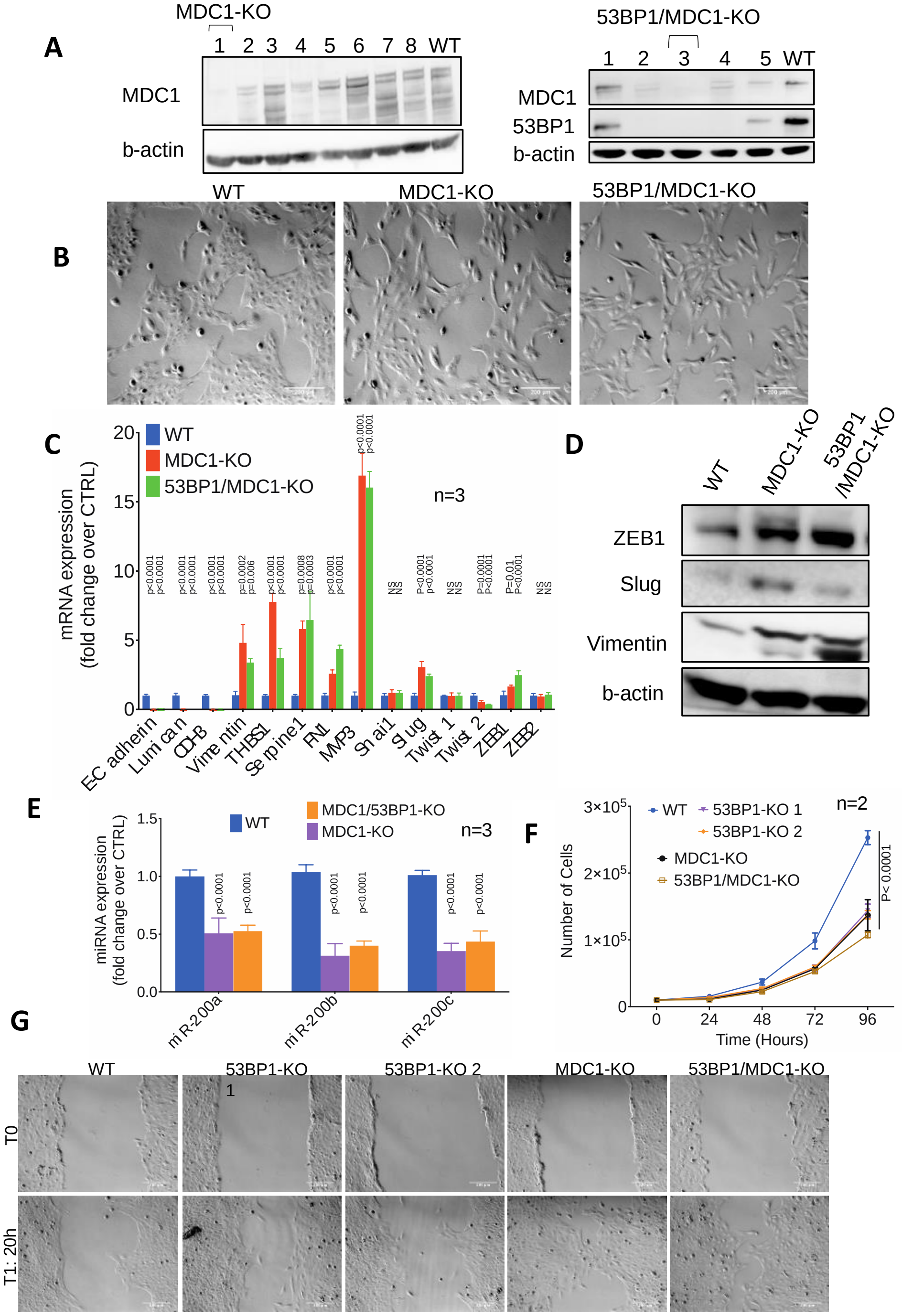


**Fig. S2.**

**Disabling the DNA Damage Response induces EMT.**

**A**. Identification by western blot of CRISPR-Cas9 transfected HEK-Early clones full KO for either DMC1 or both MDC1 and 53BP1. **B**. Cell morphology of KO clones compared with the parental cell line (WT). **C**. Relative mRNA level of expression of epithelial and EMT markers in MDC1- or 53BP1/MDC1-DKO clones compared to the parental cell line (WT). p values from paired comparisons are indicated. NS: non-significant. **D**. Western blot analyses for EMT-related markers in MDC1- and 53BP1/MDC1-DKO clones. **E**. Relative expression level of three members of the miR-200 family upon deletion of MDC1 or of both 53BP1 and MDC1. p values from paired comparisons are indicated. **F**. Growth curves for WT and 53BP1-, MDC1- and 53BP1/MDC1-DKO clones. **G**. Representative images at 0h and 20h used for the wound healing assays carried out in 53BP1-, MDC1- and 53BP1/MDC1 double KO. Quantifications are presented in Figure 1J.

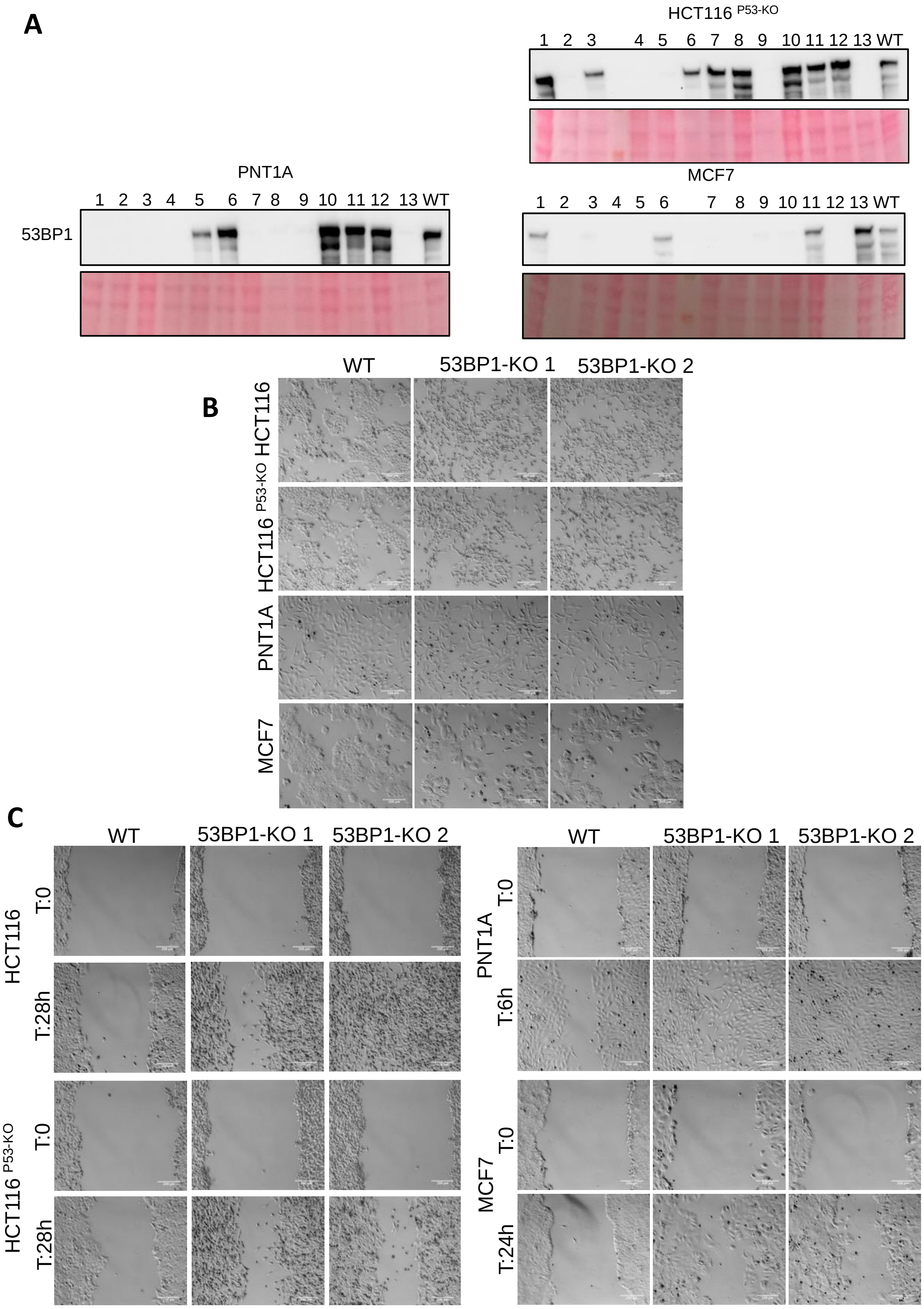


**Fig. S3.**

**EMT induction following DDR attenuation is a widespread phenomenon in cancer cells.**

**A**. Identification by western blot of CRISPR-Cas9 transfected HCT116, HCT116$^{P53\text{-}KO}$, MCF7 and PNT1A clones full KO for 53BP1. **B**. Cell morphology aspects of different 53BP1-KO clones compared with the parental cell line (WT). **C**. Representative images used for the wound healing assays carried out in 53BP1-KO clones obtained from HCT116, HCT116$^{P53\text{-}KO}$, MCF7 and PNT1A cells. Indicated readout times were determined depending on would healing capacities of parental (WT) cells. Quantifications are presented in Figure 2G.

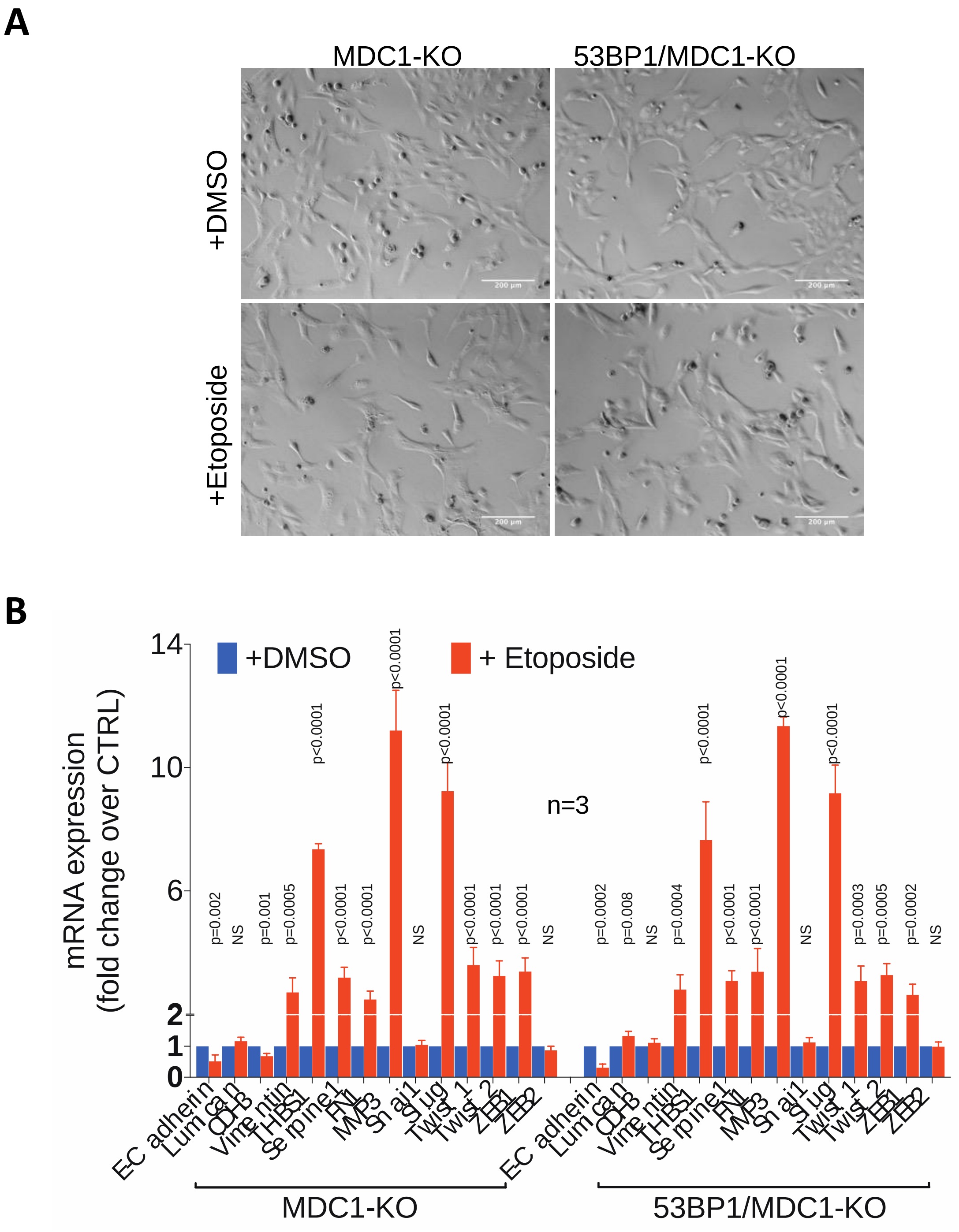


**Fig. S4.**

**Exposure to exogenous DNA damage reinforces EMT hallmarks in a DDR deficient context. A.** Cell morphology aspect of WT parental cell line and 53BP1 KO clones after treatment with etoposide 0.1 µM for 7 days. **B**. Relative mRNA level of expression of epithelial and EMT markers in MDC1- and 53BP1/MDC1-DKO clones after exposure to etoposide (0.1 µM for 7 days). p values from paired comparisons are indicated. NS: non-significant.

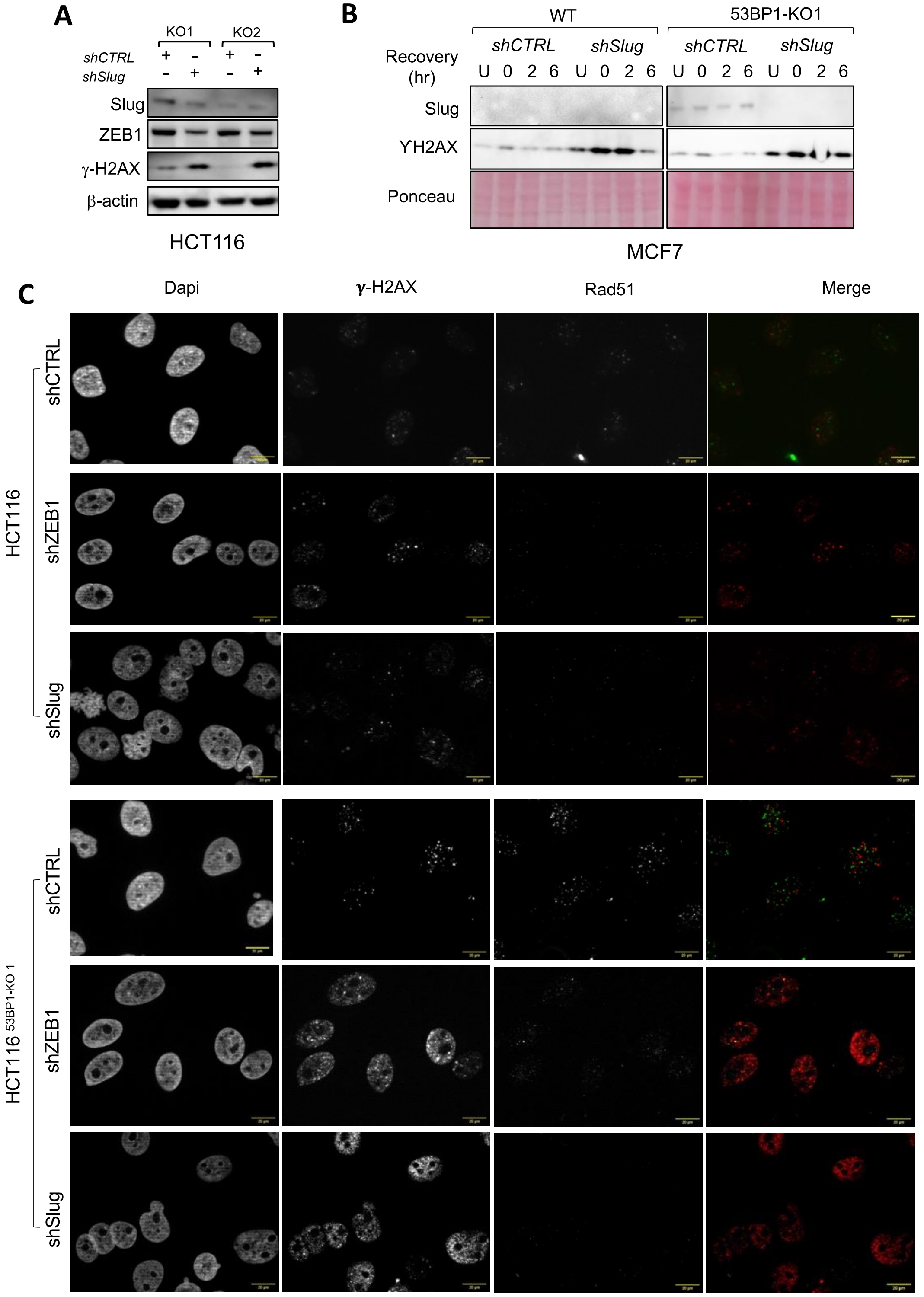


**Fig. S5.**

**EMT favors DNA repair efficiency.**

**A.** Western blot analyses for Slug and ZEB1 after introduction of an shSlug expressing vector into HCT116 53BP1-KO clones. **B**. DNA damage recovery assay in MCF7 cells, WT and KO for 53BP1, and depleted or not for Slug. Cells were treated (or not, U) with etoposide (2 μM for 24h) before medium change (0 hr) and let to recover for either 2 or 6 hours. Shown here are western blot analyses for Slug and the DNA damage marker γH2AX. **C**. Representative images immunofluorescence experiments aimed at detecting γH2AX and RAD51 foci in WT or 53BP1-KO HCT116 cells expressing either shZEB1 or shSlug. Quantitative analyses are presented in Figure 4G.

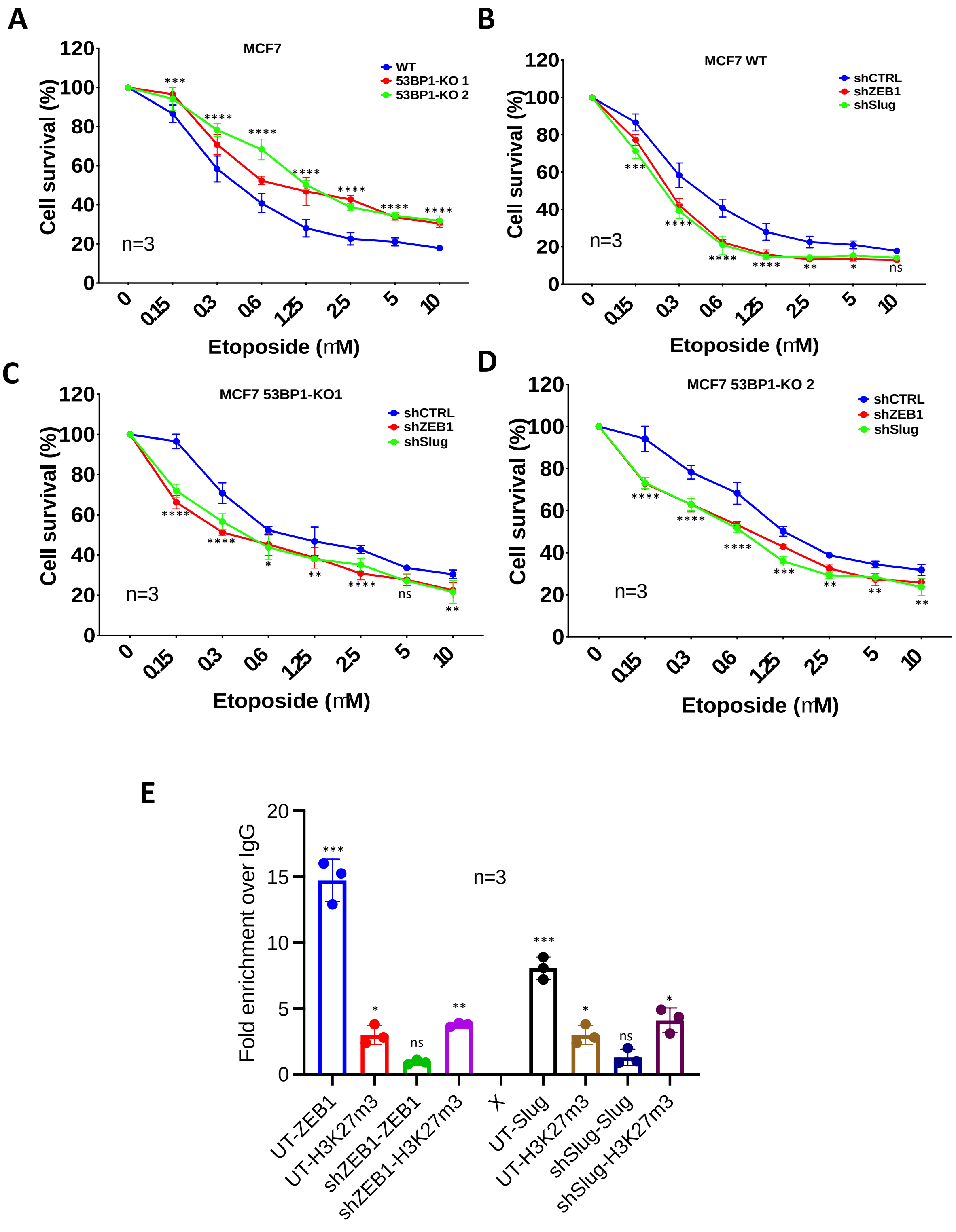


**Fig. S6.**

**EMT promotes resistance to etoposide and stimulates RAD51 expression.**

**A to D.** Cytotoxicity assays for WT and 53BP1-KO MCF7 cells, depleted or not for either ZEB1 or Slug using etoposide. Significant differences from multiple t-tests (adjusted) are shown for the following comparisons: KO vs WT (A) or shZEB1/shSlug vs shControl (B-D). **E.** Cut&Run analysis for EMT-TFs binding to the promoter of the Vimentin gene.

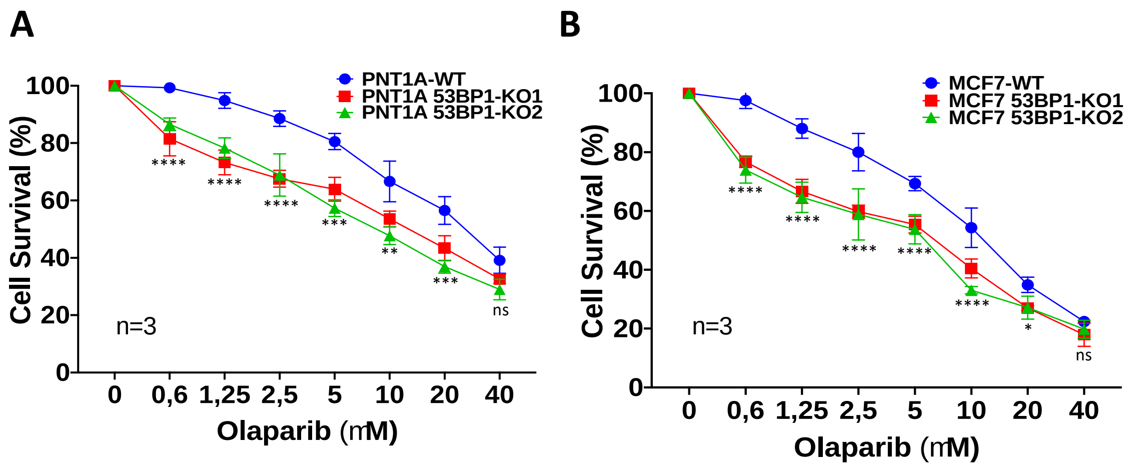


**Fig. S7.**

**EMT favors resistance against DNA damage.**

**A-B**. Cytotoxicity assays for WT and 53BP1-KO MCF7 cells as well as WT and 53BP1-KO PNT1A cells using Olaparib. Significant differences from multiple t-tests (adjusted) are shown for the comparisons KO vs WT.

**Table S1.**

Sequences of sgRNAs targeting sense and anti-sense strands in MDC1 and 53BP1 genes.

| | | |
|---|---|---|
| MDC1_S_F | accgCAGGATCAGAGCTGCTGAGA | Exon 9 |
| MDC1_S_R | aaacTCTCAGCAGCTCTGATCCTG | Exon 9 |
| MDC1_AS_F | accgCAGGTGGCATCTTGCAATTC | Exon 9 |
| MDC1_AS_R | aaacGAATTGCAAGATGCCACCTG | Exon 9 |
| 53BP1_S_F | accgGCAGTCCCACAGAGCAAGAA | Exon 10 |
| 53BP1_S_R | aaacTTCTTGCTCTGTGGGACTGC | Exon 10 |
| 53BP1_AS_F | accgGAACGATAAAAGGAGTAGAT | Exon 10 |
| 53BP1_AS_R | aaacATCTACTCCTTTTATCGTTC | Exon 10 |

**Table S2.**

Sequences of forward and reverse primers bracketing CRISPR targeted sequences.

| | | | |
|---|---|---|---|
| MDC1-F | CTGCTTGGAACTCAGCCACC | 20 | GENOTYPIC SCREENING |
| MDC1-R | GAAGATATAGAGATGACTTGTGGAATAGGAGG | 32 | GENOTYPIC SCREENING |
| 53BP1-F | AAGGAATTCTTCAGATCTTGTTGC | 24 | GENOTYPIC SCREENING |
| 53BP1-R | CAAGGCAGAAAAAGTGTTGCTC | 22 | GENOTYPIC SCREENING |
| MDC1-F | CCAAAGTGATCCTGGAGAGAGATAC | 25 | DNA SEQUENCING |
| MDC1-R | CTCTCCTCCATTAGACTGGGATCTA | 25 | DNA SEQUENCING |
| 53BP1-F | TATTTCCTAGCACTGCTCATTTTGC | 25 | DNA SEQUENCING |
| 53BP1-R | CTGAAGGGCTCCTCAAGTGC | 20 | DNA SEQUENCING |

**Table S3.**

Antibodies used for immunoblots analyses.

| Target | Host | Dilution ratio | Supplier | Reference number |
|---|---|---|---|---|
| **γ**-H2AX | Rabbit | 1:1000 | Cell Signaling | 9718 |
| H2AX | Rabbit | 1:1000 | abcam | ab11175 |
| B-Actin | HRP | 1:10000 | Santa Cruz | sc-47778 |
| 53BP1 | Rabbit | 1:5000 | novusbio | NB100-304 |
| MDC1 | Rabbit | 1:5000 | abcam | ab11171 |
| Vimentin | Rabbit | 1:1000 | Cell signaling | 9782 |
| Slug | Rabbit | 1:1000 | Cell signaling | 9782 |
| Snai1 | Rabbit | 1:1000 | Cell signaling | 9782 |
| ZEB1 | Mouse | 1:2000 | Origene | TA802298 |
| ZEB1 | Rabbit | 1:1000 | Cell signaling | 9782 |
| ZO-1 | Rabbit | 1:1000 | Cell signaling | 9782 |
| E-Cadherin | Rabbit | 1:1000 | Cell signaling | 9782 |
| Claudin-1 | Rabbit | 1:2000 | Cell signaling | 9782 |
| Twist1 | Mouse | 1:1000 | Santa Cruz | sc-81417 |
| Rad51 | Rabbit | 1:1000 | Sigma-Aldrich | PC130 |
| P-Kap1 (S824) | Rabbit | 1:1000 | Bethyl | A300-767A-M |
| Kap1 (TIF1β) | Rabbit | 1:1000 | Cell signaling | 4124 |
| Phospho-ATM | Rabbit | 1:10,000 | abcam | ab81292 |
| ATM | Rabbit | 1:10,000 | abcam | ab32420 |

**Table S4.**
Sequences of forward and reverse oligos used for RT-qPCR.

| | |
|---|---|
| FN1-F | ACAACACCGAGGTGACTGAGAC |
| FN1-R | GGACACAACGATGCTTCCTGAG |
| THBS1-F | GCTGGAAATGTGGTGCTTGTCC |
| THBS1-R | CTCCATTGTGGTTGAAGCAGGC |
| Serpine1-F | CTCATCAGCCACTGGAAAGGCA |
| Serpine1-R | GACTCGTGAAGTCAGCCTGAAAC |
| B-Actin-F | CACCATTGGCAATGAGCGGTTC |
| B-Actin-R | AGGTCTTTGCGGATGTCCACGT |
| MMP3-F | CACTCACAGACCTGACTCGGTT |
| MMP3-R | AAGCAGGATCACAGTTGGCTGG |
| twist1-F | GCCAGGTACATCGACTTCCTCT |
| twist1-R | TCCATCCTCCAGACCGAGAAGG |
| twist2-F | GCAAGATCCAGACGCTCAAGCT |
| twist2-R | ACACGGAGAAGGCGTAGCTGAG |
| ZEB1-F | GGCATACACCTACTCAACTACGG |
| ZEB1-R | TGGGCGGTGTAGAATCAGAGTC |
| ZEB2-F | AATGCACAGAGTGTGGCAAGGC |
| ZEB2-R | CTGCTGATGTGCGAACTGTAGG |
| CDH3-F | CAGGTGCTGAACATCACGGACA |
| CDH3-R | CTTCAGGGACAAGACCACTGTG |
| Vimentin-F | CGAGGACGAGGAGAGCAGGATTTCTC |
| Vimentin-R | GGTATCAACCAGAGGGAGTGA |
| E-Cadherin-F | GCCTCCTGAAAAGAGAGTGGAAG |
| E-Cadherin-R | TGGCAGTGTCTCTCCAAATCCG |
| SNAIL1-F | ACCACTATGCCGCGCTCTT |
| SNAIL1-R | GGTCGTAGGGCTGCTGGAA |

| | |
|---|---|
| SNAIL2-F | TGTTGCAGTGAGGGCAAGAA |
| SNAIL2-R | GACCCTGGTTGCTTCAAGGA |
| Rad51-Promoter-1-F | AATGTCTTCCACTTCGCCC |
| Rad51-Promoter-1-R | TTCACGCCAGTAATCCCAG |
| Rad51-Promoter-2-F | CCATTTCCCACTTCTATCCATC |
| Rad51-Promoter-2-R | GTTGCCGTCTTCTGTTTACC |
| Alu-F | TACAAAAAATTAGCCGGGCG |
| Alu-R | GATCTCGGCTCACTGCAAG |
